# A histidine cluster determines YY1-compartmentalized coactivators and chromatin elements in phase-separated super-enhancers

**DOI:** 10.1101/2021.09.15.460559

**Authors:** Wenmeng Wang, Shiyao Qiao, Guangyue Li, Cuicui Yang, Chen Zhong, Daniel B. Stovall, Jinming Shi, Dangdang Li, Guangchao Sui

## Abstract

As an oncogenic transcription factor, Yin Yang 1 (YY1) regulates enhancer and promoter connection. However, gaps still exist in understanding how YY1 coordinates coactivators and chromatin elements to assemble super-enhancers. Here, we demonstrate that YY1 activates FOXM1 gene expression through forming liquid-liquid phase separation to compartmentalize both coactivators and enhancer elements. In the transactivation domain of YY1, a histidine cluster is essential for its activities of forming phase separation, which can be extended to additional proteins. Coactivators EP300, BRD4, MED1 and active RNA polymerase II are components of YY1-rich nuclear puncta. Consistently, histone markers for gene activation, but not repression, colocalize with YY1. Importantly, multiple enhancer elements and the FOXM1 promoter are bridged by YY1 to form super-enhancers. These studies propose that YY1 is a general transcriptional activator, and promotes phase separation with incorporation of major coactivators and stabilization by distal enhancers to activate target gene expression.

## INTRODUCTION

Gene expression mediated by RNA polymerase II (Pol II) is a complex but well-regulated biological process. Transcription factors (TFs) and cofactors constitute the transcription machinery that recognizes specific elements on target promoters and coordinates concerted Pol II action (1,2). Among hundreds of TFs, many master TFs can establish super-enhancers through associating with coactivators, such as mediator and BRD4, to activate key cell identity genes (3). During malignant transformation, deregulated TFs, especially in their increasingly expressed situations, promotes oncogenic signaling pathways and processes, and thus serves as both key regulators of cancer development and promising therapeutic targets (4,5). A TF protein generally comprises a DNA-binding domain(s) (DBD) and an activation domain(s) (AD). Many DBDs have been well-classified to have conserved sequences and structural motifs shared by different families of TFs, but structural features of ADs are much less characterized (6,7). Recent studies revealed that multiple molecules of a master TF utilize its DBD to bind a target promoter and enhancers, and simultaneously use the AD to recruit various coactivators, leading to the formation of a super-enhancer complex in spatial vicinity of transcription start sites (TSS) of a target gene. A TF capable of using this mechanism typically contains intrinsically disordered regions (IDRs) in its AD(s), which allows it to form phase-separated droplets (8-13).

YY1 was first identified as a negative regulator of p53 by us and others in 2004 (14,15). Since then, the proliferative role of YY1 in oncogenesis has been frequently reported, and its increasing expression in tumor cells and tissues has been demonstrated in most cancer types (16). As a master TF, YY1 regulates many cancer-associated genes through various mechanisms that have not been completely delineated (17). YY1 promotes the expression of many key TFs with oncogenic activities, such as MYC and SNAIL1 (18-21). Stimulation of these TFs may build positive feedbacks to augment oncogenic signals in cancer cells.

A number of master TFs can form super-enhancer complexes that comprise multiple TF molecules to compartmentalize several enhancer elements and many coactivator molecules to relatively separated condensates, leading to utmost activation of target genes (8,9,22). This type of regulation requires DBDs of TFs to bind target promoters and different enhancer elements, and ADs to form phase-separated condensates with coactivators (8,10). The ADs of master TFs with phase separation capability typically possess intrinsically disordered regions (IDRs) characterized by either enrichment of acidic, proline, serine, threonine, or glutamine residues in their primary sequences, or special secondary structures (8,23-26).

YY1 promiscuously interacts with numerous transcriptional coactivators, such as EP300, CBP and PRMT1 (16), and plays a pivotal role in stabilizing enhancer-promoter loops (27). However, it still remains undetermined how YY1 coordinates the coactivators and chromatin elements to assemble super-enhancer complexes. To answer this question, we interrogate whether YY1 exerts its transcriptional activity through a phase separation mechanism. Using a variety of biochemical and cell biology approaches, we discover that YY1 possesses bona fide forming phase separation ability both in vitro and in cells. The histidine cluster of YY1 is indispensable for its capability in generating phase separation and maintaining cell proliferation. YY1-rich nuclear puncta comprise major transcription coactivators and overlap with general histone markers of gene activation. With the forkhead box protein M1 (FOXM1) as an exemplary target gene, we demonstrate that YY1 compartmentalizes coactivators and enhancer elements in phase-separated condensates to activate gene expression.

## MATERIAL AND METHODS

### Reagents, antibodies and plasmids

Reagents and antibodies used in this study include: YY1 (H-10) (Santa Cruz, cat#sc-7341, 1:1000 for Western blot (WB), 1:300 for immunofluorescent staining (IF)); YY1 (H414) (Santa Cruz, cat#sc-1703, 1:50 for chromatin immunoprecipitation (ChIP)); FOXM1 (Thermo Fisher Scientific, cat#702664, 1:5000 for WB); Flag (Sigma, cat#F1804, 1:300 for IF); EP300 (Cell Signaling Technology, cat#86377, 1:500 for IF, 1:50 for (ChIP); BRD4 (Santa Cruz, cat# sc-518021, 1:200 for IF, 1:50 for ChIP); MED1 (Santa Cruz, cat#sc-74475, 1:200 for IF, 1:50 for ChIP); CDK9 (Cell Signaling Technology, cat#2316, 1:100 for IF); RNA Pol II-S2P (Millipore, cat#04-1571, 1:200 for IF); RNA Pol II-S5P (Millipore, cat#04- 1572, 1:200 for IF); H3K4me1 (Cell Signaling Technology, cat#5326, 1:500 for IF); H3K4me3 (Cell Signaling Technology, cat#9751, 1:200 for IF); H3K27ac (Abcam, cat#ab4729, 1:500 for IF); H3K9me3 (Cell Signaling Technology, cat#13969, 1:500 for IF); pAKT-Thr308 (Cell Signaling Technology, cat#13038S, 1:1000 for WB); pAKT-Ser473 (Cell Signaling Technology, cat#4060S, 1:1000 for WB); AKT (Cell Signaling Technology, cat#4685S, 1:1000 for WB); GAPDH (Acton, cat#10R-G109A, 1:1000 for WB); Goat anti-Rabbit IgG (H+L) Highly Cross-Adsorbed Secondary Antibody, Alexa Fluor 488 (Thermo Fisher Scientific, cat#A32731, 1:500 for IF); Goat anti-Mouse IgG (H+L) Cross-Adsorbed Secondary Antibody, Alexa Fluor 594 (Thermo Fisher Scientific, cat#A32742, 1:500 for IF); Goat anti-mouse IgG-HRP (Santa Cruz, cat#sc-2005,1:5000 for WB); Goat anti-rabbit IgG-HRP (Santa Cruz, cat#sc-2004, 1:5000 for WB); JQ-1 carboxylic acid (MedChemExpress, cat#202592-23-2); 1,6-hexanediol (Aladdin, cat#H103708).

The full-length coding sequences of YY1, HOXA1, FOXG1B, ZIC3 and HNF6 (or their variants), as well as IDRs of EP300, BRD4, MED1 and RNA Pol II, were individually subcloned into a modified version of a pGEX vector with 6×His and EGFP or mCherry at the N-terminus. A bacterial expression system was used to express these coding sequences and recombinant proteins were purified using Ni-NTA agarose. Meanwhile, the full lengths of YY1 or its mutants, and EP300 were subcloned into a eukaryotic EGFP or mCherry expression vector. YY1 or its mutants, and FOXM1 coding sequences were also cloned into a lentiviral vector with a 3×Flag-tag at the N-terminus. An shRNA, sh-YY1, targeting the 3’-UTR of YY1 mRNA, and a control shRNA (sh-Cont) were designed as previously described (28). The promoter of FOXM1 was amplified by a nest PCR method and subcloned upstream of the Gaussia luciferase (Gluc) coding sequence to generate a pFOXM1-prmt-Gluc vector (WT). Also, reporter constructs S1M, S2M, S3M, S4M, S5M, S4/5M, S1/3M and S2/4/5M with correspondingly mutated YY1 binding sites were generated using the ClonExpress^®^ II Recombination system (Vazyme Biotech Co., Ltd., Nanjing, China). The predicted enhancer regions were also individually amplified by the nest PCR method and subcloned upstream of the FOXM1 promoter or downstream of Gluc to generate reporter constructs.

### Protein expression and purification

All His⨯6-tagged constructs were overexpressed in *E. coli* BL21 (DE3) cells. Bacteria were cultured grown to an optical density at 600 nm (OD_600nm_) of 0.6, and induced overnight with 0.15 mM IPTG at 18°C. Bacteria were pelleted and resuspended in a lysis buffer (20 mM HEPES, 0.2 mM EDTA, 100 mM KCl, 20% Glycerol, 1% Triton, 2 mM PMSF, 1 mg/mL lysozyme). After sonication, the bacterial lysate was centrifuged at 12000 g for 30 min. His⨯6-tagged proteins in the supernatant were purified by Ni-NTA agarose beads (GE Healthcare). After extensive wash by a buffer containing 20 mM imidazole, the fractions eluted by 400 mM imidazole were collected and dialyzed. The size and purity of the purified proteins were monitored by SDS-PAGE.

### Cell culture, transfection, lentiviral production, and infection

The HeLa, HEK-293T, U2OS, MDA-MB-231, MCF-7 and MCF-10A cells were cultured according to the protocols of the ATCC. All culture media were purchased from Gibco and fetal bovine serum (FBS) was from ExCell Bio. Transfection of cells was carried out using Lipofectamin 2000 (Thermo Fisher Scientific, Shanghai, China) according to the manufacturer’s instructions. Lentiviral production and infection followed our published procedure (29).

### *In vitro* phase separation assay

Recombinant EGFP- or mCherry-fusion proteins were diluted to appropriate concentrations using 50 mM Tris·HCl, pH 7.4. Recombinant proteins were added to solutions containing 125 mM NaCl and 10% PEG-8000 as crowding agents unless specified. Protein solution (5 μl) was immediately loaded onto a glass slide, and covered with a coverslip. Slides were then imaged with a ×60 objective using the GE Dela Vision Elite (GE, Boston, MA, USA).

Turbidity experiments were performed in tubes. Samples (60 μl) containing appropriate concentrations of proteins, NaCl and 10% PEGs with indicated molecular weights were left to stand for 10 s at room temperature. OD_600nm_ was measured using BioSpectrometer basic (Eppendorf, Hamburg, Germany).

### Fluorescence recovery after photo-bleaching (FRAP) imaging

FRAP was performed on a Zeiss LSM880 microscope using a ×63 oil-immersion objective. Images were acquired using the ZEN software. FRAP of the central region of protein droplets, three iterations of bleaching were performed with a 488 nm Argon laser at a 100% power with 3 frames being acquired prior to the bleach pulse. Fluorescence recovery was recorded every 2 s for 400 s after bleaching. U2OS cells, which were transfected for 24 h and cultured in glass-bottom dishes (NEST, China), were analyzed using FRAP studies. Three iterations of bleaching were performed with a 488 nm Argon laser at 30% power. Fluorescence recovery was recorded every 0.8 s for 20 s after bleaching. Analyses of the fluorescence intensity of bleached region, reference region and background region were carried out using the FRAP module in the ZEN software.

### Immunofluorescence staining and live-cell imaging

Cells were seeded on glass coverslips in 12-well plates and cultured overnight. Subsequently, cells were fixed and permeabilized with Immunol Staining Fix Solution (Beyotime) for 30 min at room temperature. After blocked by 10% FBS for 30 min at room temperature, cells were incubated primary antibody for 30 min at room temperature. Then, the cells were washed thrice with PBS and incubated with Alex-Fluor-488 or 594-conjugated secondary antibodies (Thermo Fisher Scientific, Shanghai, China) for 30 min at RT. Finally, the cells were washed thrice with PBS and nuclei were counterstained with DAPI (Beyotime), and images were captured by the GE Dela Vision Elite (GE, Boston, MA, USA).

For live-cell imaging, U2OS cells were seeded on glass-bottom dishes and transfected by EGFP or mCherry fusion plasmids. After 24 h of transfection, nuclei were counterstained with Hoechst (Beyotime), and cells were imaged using the GE Dela Vision Elite (GE, Boston, MA, USA) with a ×60 objective. During image acquisition, cells were incubated in a chamber at 37 °C with 5% CO_2_.

### Western blot analysis

Cells were washed and lysed in a protein lysate buffer. Total protein concentrations were measured using the Bradford protein method. Protein samples were separated on the SDS-PAGE and transferred into poly-vinylidene difluoride transfer (PVDF) membranes. The membranes were blocked by 5% nonfat milk in TBST buffer at room temperature for 1 h, and incubated with primary antibodies in TBST buffer for 4°C overnight. After three washes with TBST, the membranes were incubated with corresponding secondary antibodies at room temperature for 1 h. The membranes were washed and then visualized using ECL kit (Vazyme Biotech Co., Ltd., Nanjing, China).

### Reverse transcription and quantitative PCR (RT-qPCR)

Total RNAs were extracted from cultured cells using the TRIzol reagent (Thermo Fisher Scientific Inc., Shanghai, China), and cDNA was synthesized using M-MLV reverse-transcriptase (Vazyme Biotech Co., Ltd., Nanjing, China). In the reaction of reverse transcription, 1 μg of total RNA was mixed with 0.5 μg/μl of oligo(dT) primer, followed by incubation at 65°C for 5 min and 4°C for 2 min. The tubes were then immediately incubated at 42°C for 30 min and chilled at 4°C. For qPCR, cDNA was amplified using gene specific primers and the LightCycler 480 SYBR Green PCR Master Mix (Roche, Basel, Switzerland) on Lightcycler 480 instrument (Roche). The conditions used for qPCR were as follows: 95°C for 3 min, followed by 40 cycles of 95°C for 15 s and 60°C for 1 min. All reactions were performed in triplicate. The results were analyzed using the 2^-ΔΔCt^ method and normalized using β- actin. Primer sequences for qPCR were as follows: β-actin (5′-TTCCTTCCTGGGCATGGAGT and 5′- TCTTCATTGTGCTGGGTGCC), YY1 (5′- CCCACGGTCCCAGAGTCCA and 5′- GTGTGCGCAAATTGAAGTCC), and FOXM1 (5′-GCAGGCTGCACTATCAACAA and 5′- TCGAAGGCTCCTCAACCTTA).

### Chromatin immunoprecipitation (ChIP)

ChIP analysis was performed as previously reported (30). In brief, cross-linking was completed after cell culture, followed by nuclei preparation and chromatin digestion. DNA gel electrophoresis was used to confirm adequate digestion. Samples immunoprecipitated by a normal IgG and a specific antibody were purified and subjected to semiquantitative both PCR and qPCR. For semiquantitative PCR, PCR products were analyzed using gel electrophoresis. The sequences of primers were listed in Supplementary Table S1.

### Electrophoretic mobility shift assay (EMSA)

Oligonucleotides labelled with Cy5 at the 5’-end were synthesized and purified using HPLC method by Genewiz (Suzhou, China), and sequences of oligonucleotide probes and competitors were listed in Supplementary Table S2. The EMSA was conducted as we previously described (31). Briefly, 1 μg of a purified His×6-tagged YY1 protein was incubated with 0.5 pmol of labeled double-stranded probe in a binding buffer (250 mM HEPES, 500 mM KCl, 20 mM MgSO_4_, and 10 mM DTT, pH 8.0) on ice for 30 min. In the competitive binding experiments, excessive unlabeled probes were added to the binding reaction of the labeled probe and His⨯6-YY1. After the binding reaction, the samples were separated by 8% native PAGE at 100 V for 50 min at 4°C. The fluorescent intensity of the bands was immediately determined by Typhoon FLA7000 (GE, Boston, MA, USA).

### Luciferase reporter assay

The reporter plasmids were constructed as described above. Cells in 24-well plates were cotransfected by 250 ng of a reporter construct, 500 ng of shRNA or expression construct, and 20 ng of pCMV-SEAP (secreted alkaline phosphatase) as a control. The Gluc activity was measured at 48 h after transfection as described previously (31).

### Chromosome Conformation Capture (3C) experiments

The 3C analysis was performed following previously described procedures with minor modifications (32). Total of 1⨯10^6^ MDA-MB-231 or MCF-7 cells were cross-linked by 2% formaldehyde in 10% (v/v) FBS/PBS for 10 min at room temperature, and the reaction was stopped by adding glycine to a final concentration of 0.125 M. After washed by PBS, the fixed cells were incubated for 15 min in cold lysis buffer (10 mM Tris·HCl, pH 8.0, 10 mM NaCl, 0.2% NP-40) with 1×complete protease inhibitor cocktail (Roche). The nuclei were harvested and resuspended in a restriction buffer (50 μl 10×CutSmart buffer and 0.3% SDS), and incubated for 1 h at 37°C. After incubated with 1.8% Triton X-100 for 1 h at 37°C to sequester the SDS, 600 units of restriction enzyme EcoRI or HindIII (NEB) was used to digest genomic DNAs by incubating at 37°C overnight. Then, 1.6% SDS was used to inactivate the restriction enzyme by incubation at 65°C for 20 min. The solution was diluted by adding 7 ml of ligation buffer (NEB) containing 1% Triton X-100 and 30 Units of T4 DNA ligase (NEB), followed by ligation reaction at 16°C for 4.5 h and then at room temperature for 30 min. Cross-linking was released by Proteinase K digestion at 65°C for 16 h. Finally, DNA fragments were purified by phenol-chloroform extraction and ethanol precipitation. The ligation products were analyzed by PCR using primers located adjacent to EcoRI or HindIII digested sites. PCR products were cloned into a pBluescript plasmid and sequenced to verify the ligated fragments. The sequences of primers used in 3C assay are shown in Supplementary Tables S3 and S4.

### Cell viability, colony formation, and wound healing assay

These assays were performed as previously reported (33). In these experiments, cells were infected by lentivirus expressing either shRNAs or cDNAs. Each experiment was performed in triplicate.

### Statistical analysis

All data were derived from at least three independent experiments. Results were presented as a mean with either standard deviation (SD) or standard error of mean (SEM), and sample numbers are indicated unless otherwise noted in the figure legends. Statistical significance calculations comparing two conditions were performed using a two-tailed unpaired Student’s *t-test*. The criterion of statistical significance level was denoted as follows: \**P* < 0.05; \*\**P* < 0.01; \*\*\**P* < 0.001.

## RESULTS

### YY1 undergoes liquid-liquid phase separation

YY1 has a highly acidic transactivation domain (TAD) at N-terminus; its N-terminal 154 amino acids contain 76 negatively charged glutamic/aspartic acid (E/D) residues, but only two positively charged residues (R109 and R122), as well as 18 histidines (Hs) and 20 glycines (Gs) (Supplementary Figure S1A). To date, structure of full length YY1 has not been reported, although cocrystal structure of its C-terminus with a binding element was resolved (34). YY1 TAD embraces an eleven-E/D (11×E/D) cluster and an 11×H cluster flanked by two acidic regions (Figure 1A, top row). Consistently, inspection of YY1 primary sequence by the IUPred and VSL2 algorithms (35,36) revealed a high propensity of structural disorder in its TAD (Figure 1A, bottom row). Amino acid composition examination revealed multiple potential IDRs in YY1, and the strongest one contains an 11×E/D and an 11×H clusters separated by a G-rich stretch (Figures 1A and 1B). Importantly, E/D and H clusters, and their proximate regions are highly conserved among different species of vertebrates (Figure 1B). Thus, bioinformatic analyses strongly support the presence of an IDR in YY1’s TAD.

**Figure 1.**
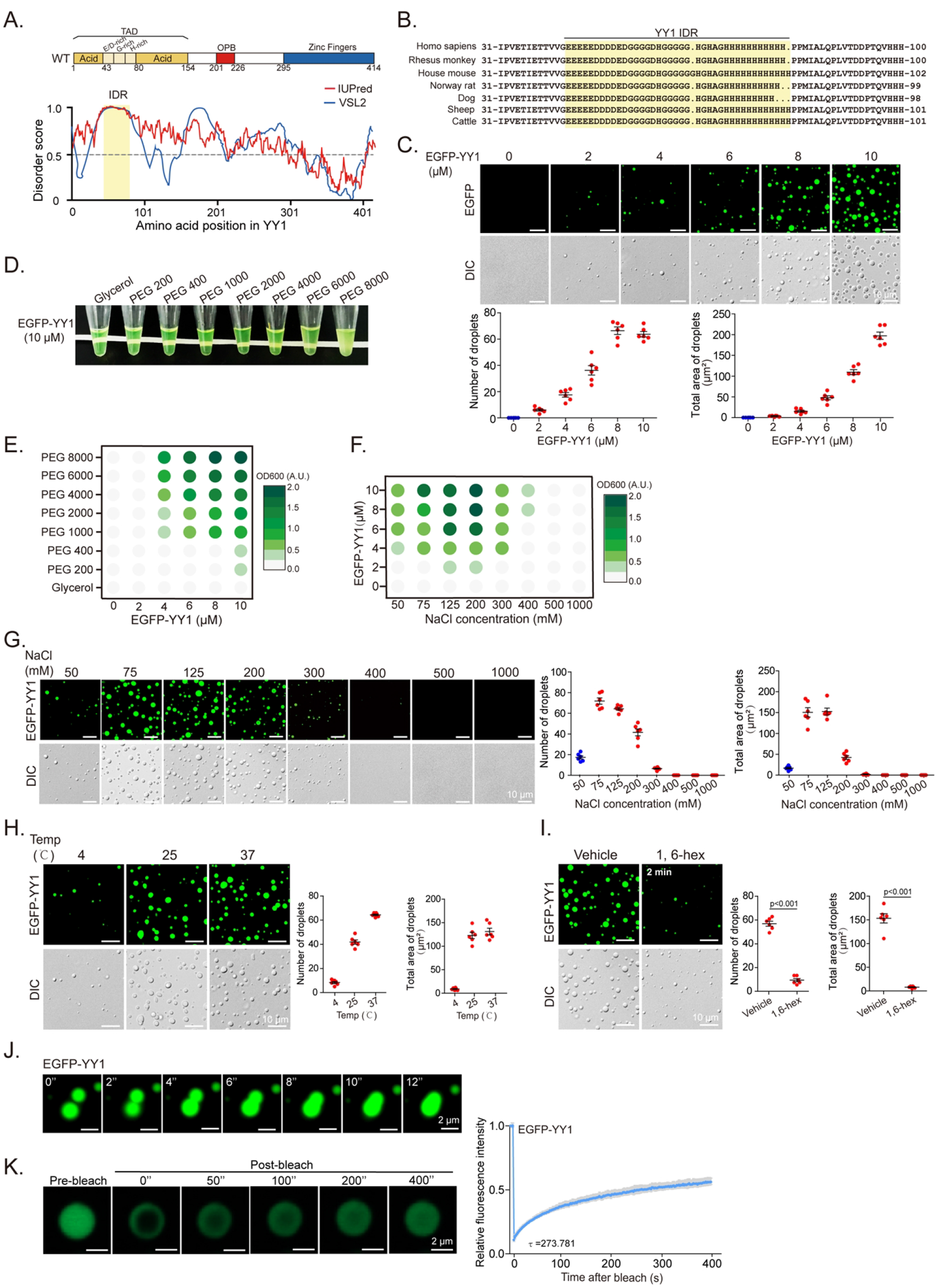
YY1 undergoes liquid-liquid phase separation (LLPS) *in vitro*. **A**. Domain structure and graphs of YY1 IDRs based on VSL2 and IUPred algorithms. Scores > 0.5 indicate disorder. Yellow shade depicts potential IDR. **B**. IDR regions of YY1 proteins from different vertebrates. **C**. Representative fluorescence and differential interference contrast (DIC) images of EGFP-YY1 droplets at different protein concentrations in a buffer containing 125 mM NaCl and 10% PEG-8000 (the same condition hereafter, if not specified). Quantification of droplets’ numbers and area is shown at bottom row. **D**. Turbidity visualization of EGFP-YY1 droplet formation. Tubes containing EGFP-YY1 (10 μM, the same concentration hereafter, if not specified) in the buffer containing PEGs with increasing molecular weights. **E** and **F**. OD_600nm_ phase diagram of turbidity changes caused by EGFP-YY1 droplet formation in different PEGs (**E**) and an increasing NaCl concentrations (**F**). **G**. Representative fluorescence and DIC images of EGFP-YY1 droplets at increasing NaCl concentrations. Quantification of droplet’s numbers and area is shown at right. **H** and **I**. EGFP-YY1 droplet formation at different temperatures (**H**), and in the presence or absence of 5% of 1,6-hexanediol (1,6-hex). Quantification is shown at right. In **C, G-I**, quantification of droplets’ numbers and area was mean ± s.e.m. from 6 fields each group. **J**. Two EGFP-YY1 droplet coalescence over a time course. **K**. Fluorescence recovery after photobleaching (FRAP) of EGFP- YY1 droplets. FRAP curve is shown at right. Data are presented as mean ± s.d. (n = 6 droplets).

Many IDR-containing TFs form dynamic liquid-like droplets, or gel-like phase-separated condensates, due to multivalent and weak interactions among IDRs (9,25,26). Fluorescent droplets and turbidity created by gel-like condensates are used to evaluate phase separation, while polyethylene glycol (PEG) mimicking crowded cellular conditions promotes this *in vitro* process (9). When purified recombinant EGFP-YY1 (Supplementary Figure S1B) was incubated with PEG of a molecular weight (MW) 8000 Dalton (PEG 8000) in the presence of 125 mM NaCl, we observed droplet formation, of which the density and size scaled up with increased protein concentrations (Figure 1C). Meanwhile, as measured by optical density at 600 nm (OD_600_), turbidity of solutions monotonically intensified, which was proportional to MWs of PEG in the buffer (Figure 1D and 1E). To evaluate biological relevance of EGFP-YY1 droplet formation, we carried out the assays in solutions with increasing salt concentrations. The highest turbidity and most droplet formation by EGFP-YY1 were observed at 125- 200 mM NaCl, in the range of physiological saline levels, but vanished as salt concentration reached 500 mM (Figures 1F, 1G and Supplementary Figure S1C). In addition, ability of EGFP-YY1 in forming droplets was higher at 37°C than that at 25°C, and markedly descended at 4°C (Figure 1H). Importantly, EGFP-YY1 droplets were sensitive to 1,6-hexanediol, a chemical disrupting liquid-liquid phase separated condensates (Figure 1I). We also observed fusion events between two adjacent droplets (Figure 1J and Supplementary Video 1) and quick green fluorescence recovery of droplets after targeted photobleaching treatment (Figure 1K), indicating a dynamic feature of EGFP-YY1 condensates.

When EGFP-YY1 was expressed in U2OS cells, we detected green fluorescent puncta that also exhibited the ability of prompt fluorescence recovery after photobleaching (Figure 2A), consistent with its droplet formation *in vitro*. Furthermore, treatment of EGFP-YY1 transfected cells by 1,6-hexanediol greatly diffused green fluorescent puncta in nuclei, suggesting their liquid condensate properties (Figure 2B). The results strongly suggest that YY1 is capable of forming phase-separated condensates. To evaluate how special amino acid clusters in the predicted IDR of YY1 (Figures 2C, left panel, and Supplementary Figure S1D) contributed to phase-separation, we generated EGFP- YY1 mutants E/D-A, G-A and H-A (with E/D-, G- and H-rich regions replaced by corresponding numbers of alanines, respectively), and their deletion mutants (denoted as Δ), as well as ΔIDR and EGFP-IDR fusion mutants (Figure 2C, left panel), and purified recombinant proteins from a bacterial expression system (Figure 2C, right panel). We tested their droplet formation capability in a droplet formation buffer containing PEG 8000 and 125 mM NaCl, and also transfected their eukaryotic vectors into U2OS cells. EGFP-IDR could steadily form droplets with comparable numbers and sizes to those of the WT, while EGFP-YY1-ΔIDR completely lost this ability (Figure 2D). Consistently, EGFP-YY1-ΔIDR did not generate nuclear puncta, but instead showed diffusive distribution in both nuclei and cytoplasm (Figure 2E). Strikingly, both H-A and ΔH mutants of EGFP-YY1 exhibited very similar phenomena to IDR-deleted mutant, indicating a critical role of 11×H cluster in YY1 phase separation. Meanwhile, E/D-A and ΔE/D mutants formed droplets with markedly reduced numbers and sizes, and displayed diffused green fluorescence in nuclei; however, G-cluster mutation or deletion had much less impact on droplet forming ability of EGFP-YY1 than that of H and D/E cluster changes, and consistently G-A and ΔG mutants could still generate puncta in cells with comparable numbers and sizes to WT (Figure 2E). The turbidity of EGFP-YY1 WT and its mutants in PEGs reflected their droplet forming capability, with the least condensates formed by H-A, ΔH and ΔIDR mutants (Supplementary Figure S1E). In addition, in transfection of U2OS cells by Flag-tagged YY1 vectors, effects of IDR alterations on punctum formation were virtually identical to that of EGFP-YY1 proteins (Supplementary Figure S1F). These data strongly suggested that both H and E/D clusters, especially the former one, are important elements of YY1’s IDR in phase separation. We also designed (7-Methoxycoumarin-4-yl) acetic acid (Mca)-labeled peptides, with a transmembrane sequence (TAT) fused to an A/G-rich control sequence, E/D and H cluster sequences of YY1 (Figure 2F). Importantly, both TAT-E/D and TAT-H formed fusion droplets with EGFP-YY1, suggesting their ability in promoting phase separation. Contribution of E/D clusters to liquid condensate formation has been frequently observed (9,24), but the role of H clusters in phase separation was only reported in P-TEFb (37). In addition to YY1, we also verified an indispensable role of H clusters to the phase separation of HOXA1, FOXG1B, ZIC3 and HNF6 proteins (Supplementary Figures S2A-S2D).

**Figure 2.**
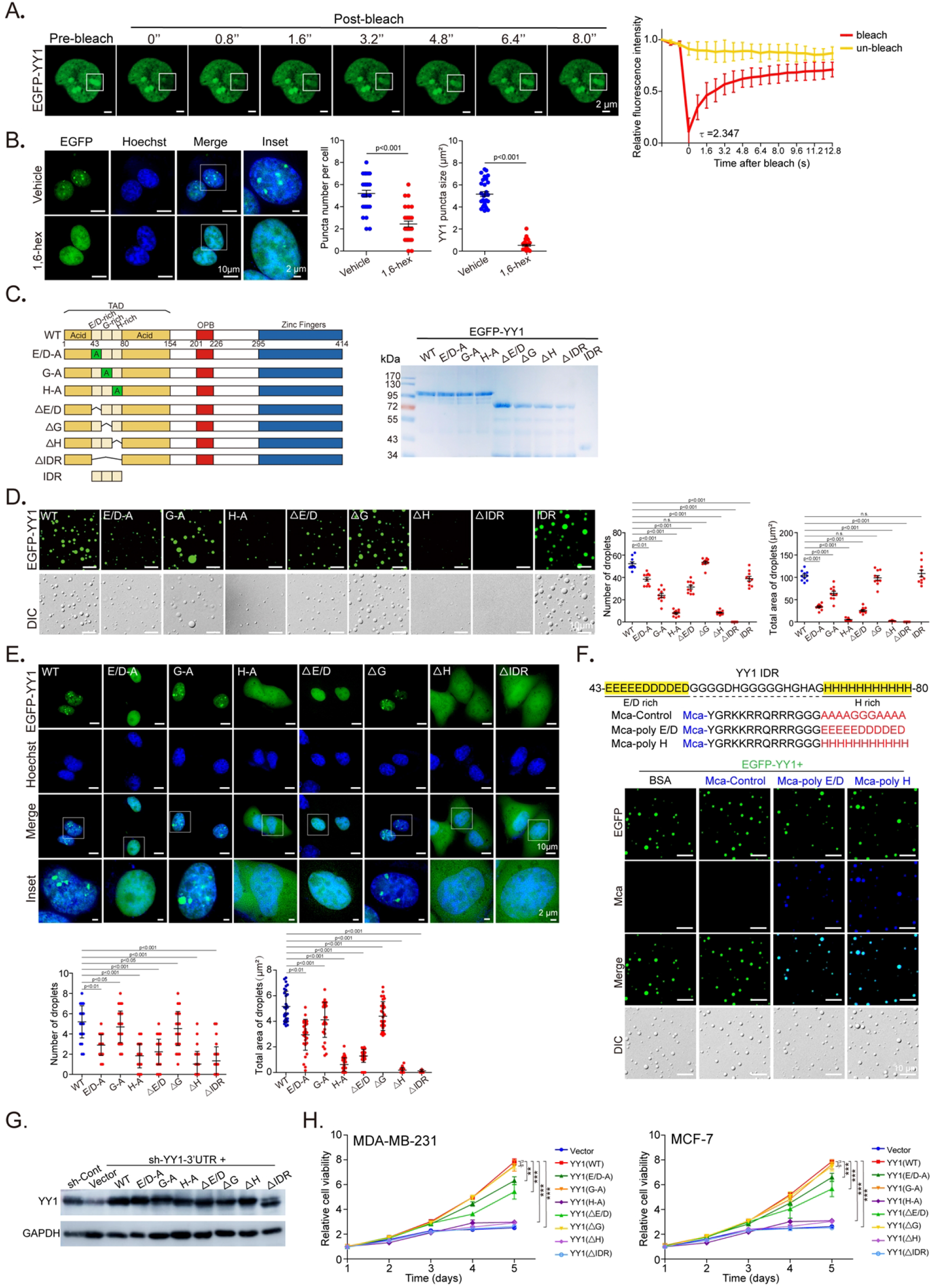
Characterization of YY1 LLPS and its functional role. **A**. FRAP of EGFP-YY1 puncta in U2OS cells. White squares depict photobleached puncta undergoing fluorescence recovery. FRAP quantification is show at right; bleaching event occurred at t = 0 s. Data represent mean ± s.d. (n = 6). **B**. EGFP-YY1 puncta in live U2OS cells treated in buffers with or without 5% 1,6-hexanediol. Nuclei were visualized by Hoechst staining. Quantification of numbers and sizes of puncta is shown at right. **C**. Domain structure of YY1 protein. E/D, G and H represent E/D-, G- and H-rich regions, respectively. SDS-PAGE of recombinant YY1 wild type (WT) and mutant proteins purified from *E. coli* is shown at right. **D**. Droplet formation of EGFP-YY1 WT and mutants. Quantification of droplets’ numbers and area is shown at right as mean ± s.e.m. (n = 6). **E**. Live U2OS cells transfected with EGFP-YY1 WT and mutants. In **B** and **E**, 30 cells were quantified for each sample. Data represent mean ± s.e.m. **F**. Representative droplet formation by EGFP-YY1 and peptides. EGFP-YY1 and (7-methoxycoumarin-4-yl) acetic acid (Mca)-labeled poly E/D, poly H and control (A/G repeat) peptides or BSA were incubated. **G**. Ectopic YY1 expression with simultaneous endogenous YY1 knockdown. MDA-MB-231 cells with DOX-induced sh-YY1 targeting 3’-UTR were infected by lentivirus of 3×Flag-YY1 WT and mutants, or a vector. Cell lysates were analyzed by Western blot using indicated antibodies. **H**. Effects of YY1 mutations on cell viability. MDA-MB-231 and MCF-7 cells expressing 3×Flag-YY1 WT and mutants with endogenous YY1 knockdown were evaluated by WST-1 assays to determine cell viability.

### Phase separation capability of YY1 is essential to its cell proliferative activity

To test whether the mutations or deletions could adversely affect YY1’s function, we expressed the mutants in MDA-MB-231 and MCF-7 cells, with simultaneous endogenous YY1 silencing by an shRNA targeting 3’-UTR of YY1 mRNA (Figure 2G). YY1 depletion reduced breast cancer cell viability, consistent with our previous report (38), while ectopically expressed WT YY1 could largely restore it (Figure 2H). In accordance to droplet and punctum formation studies, IDR deletion, H-cluster mutation or deletion completely abolished YY1’s ability in restoring cell viability, suggesting indispensability of the IDR and H-cluster to YY1’s function (Figure 2H). Similarly, both E/D-A and ΔE/D mutants only partially rescued YY1 depletion-caused viability decrease, while G-A and ΔG mutants virtually retained YY1’s function in reinstating cell viability (Figure 2H). In addition, results of YY1 mutants in promoting cell migration and clonogenicity were consistent with the data of cell viability (Supplementary Figures S2E and S2F). Overall, the IDR and H-cluster of YY1, key components for phase separation, are essential to its role in maintaining basic cellular activities.

### YY1 compartmentalizes coactivators to nuclear puncta

The name Yin Yang 1 represents its ability in mediating both repression and activation of target genes, depending on recruited cofactor (39). In the past decade, the role of YY1 in promoting gene expression has been frequently reported. Consistently, YY1 interacts with several histone acetyltransferases, including EP300, CBP and PCAF (40,41). Among them, EP300 is a well-recognized coactivator. We analyzed the primary sequence of human EP300 using the IUPred and VSL2 algorithms, and discovered five potential IDR regions, of which the IDR3 and IDR5 are larger than the others (Figure 3A). Using a bacterial expression system, we purified recombinant EGFP fusion proteins with these IDR regions (Supplementary Figure S3A). EGFP-EP300-IDR3 and -IDR5, but not the other three proteins, could generate droplets (Figure 3B), with dependencies on PEG MWs and protein concentrations (Figure 3C, Supplementary Figures S3B and S3C), and formed relatively large droplets at 25°C and 37°C versus 4°C (Figure 3D). The two EP300-IRDs formed the most droplets at NaCl concentrations between 50 and 125 mM (Supplementary Figures S3D and S3E). Droplets formed by EGFP-EP300-IDR3 and -IDR5 also showed properties of 1,6-hexanediol sensitivity (Figure 3E), adjacent droplet fusion (Figure 3F, Supplementary Videos 2 and 3), and recovery from photobleaching treatment (Figure 3G), characteristics of phase separation condensates.

**Figure 3.**
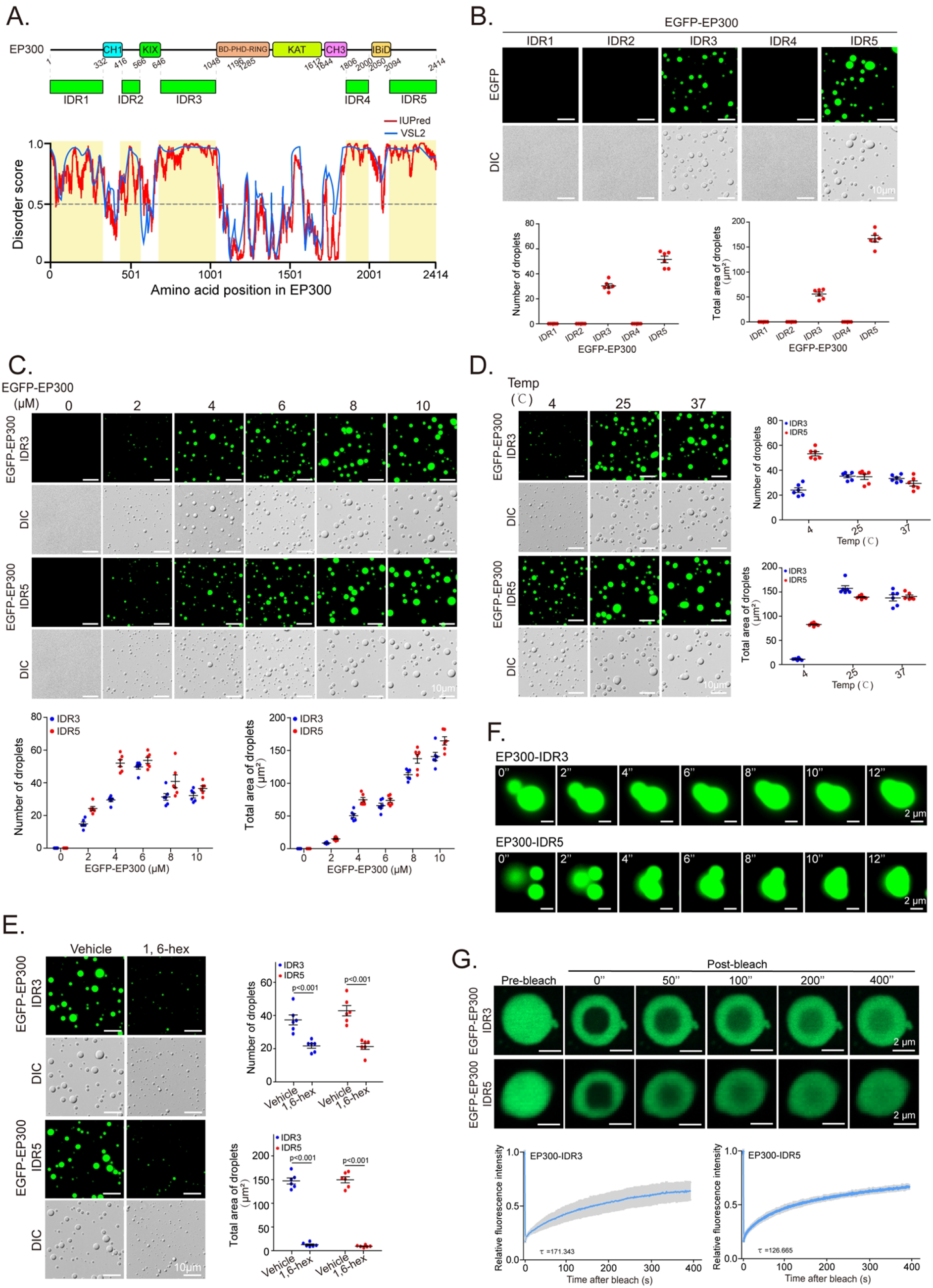
YY1 coactivator EP300 undergoes LLPS *in vitro*. **A**. EP300 domain structure and predicted IDRs (yellow shades). **B**. Droplets of EGFP-EP300-IDRs (top row) and their quantification of numbers and area (bottom row). **C**. Droplets of EGFP-EP300- IDR3 and -IDR5 at different protein concentrations. Quantification is at bottom row. **D** and **E**. EGFP- EP300-IDR3 and -IDR5 droplet formation at different temperatures (**D**), and in the presence or absence of 5% 1,6-hexanediol (**E**). In **B-E**, representative images are presented, and quantification is shown as mean ± s.e.m. from 6 droplets’ fields in each group. **F**. EGFP-EP300-IDR3 and -IDR5 droplets coalescence. **G**. FRAP recovery of EGFP-EP300-IDR3 and -IDR5 droplets. FRAP recovery curves at right are the data of mean ± s.d. (n = 6 droplets).

Consistent with *in vitro* data, immunostaining analyses also presented endogenous EP300 puncta with fluorescence recovery from photobleaching and sensitivity to 1,6-hexanediol (Figures 4A and 4B), suggesting EP300 phase separation in a cellular environment. With EGFP-EP300 and mCherry-YY1 cotransfected into U2OS cells, we observed colocalization of the green and red fluorescent signal (Figure 4C). When examining the endogenous proteins in MDA-MB-231 and MCF-7 cells, we observed that most EP300 puncta stained by its antibody were overlapped with YY1 signal (Figure 4D). Importantly, when Flag-YY1 were transfected into the cells, endogenous EP300 showed intensely stained puncta well-overlapped with Flag tag signal (Figure 4E). Importantly, puncta formed by Flag-YY1 were markedly larger in sizes than those by endogenous YY1 (Figure 4F). Consistently, mCherry-YY1 could form droplets with EGFP-EP300-IDR3 and -IDR5 with overlapped color and increased sizes, especially the latter one, compared to other predicted EP300-IDRs (Figure 4G). The data strongly suggest that YY1 is a primary TF recruiting EP300 to activate gene expression.

**Figure 4.**
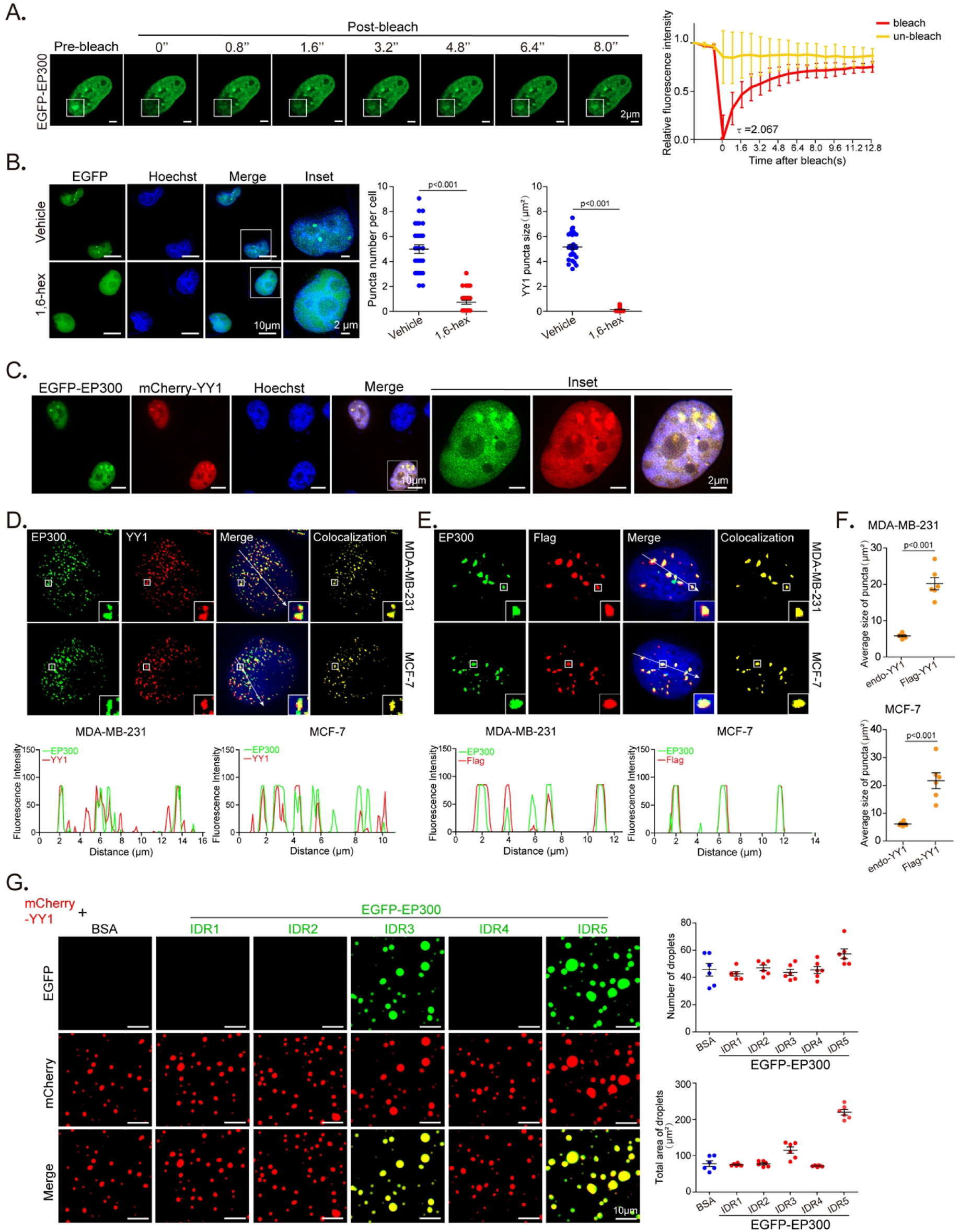
EP300 and YY1 condensates have LLPS properties. A. FRAP recovery of EGFP-EP300 puncta in U2OS cells. White squares highlight photobleached puncta, and quantification is at right. Bleaching occurs at t = 0 s. Data represent mean ± s.d. (n = 6).**B**. EGFP-EP300 puncta in live U2OS cells in the presence or absence of 5% 1,6-hexanediol. Nuclei were detected by Hoechst staining. For each sample, 30 cells were quantified. Data are presented as mean ± s.e.m. **C**. EGFP-EP300 and mCherry-YY1 puncta in live U2OS cells. Expression vectors were transfected into U2OS cells followed by fluorescent microscopy. Nuclei were detected by Hoechst staining. **D** and **E**. Colocalization of endogenous EP300 with YY1 (**D**) or Flag-YY1 (**E**) in nuclear puncta in breast cancer cells. Immunofluorescence staining was used with nuclei detected by DAPI. Line scans of colocalization images are depicted by white arrows (bottom row). **F**. Quantified average sizes of merged puncta in MDA-MB-231 (**D**) and MCF-7 (**E**) cells. Data are presented as mean ± s.e.m. with puncta in 6 fields of each group. **G**. Representative images of droplets of mCherry-YY1 incubated with EGFP-EP300-IDR mutants. Quantified droplets’ numbers and sizes in merged images is shown at right. Data are presented as mean ± s.e.m. with droplets in 6 fields for each group.

Furthermore, we tested whether YY1 cooperates with other coactivators. In immunostained MDA-MB- 231 cells, endogenous YY1 colocalized with MED1, BRD4 and CDK9 (Figure 5A). Consistently, Flag- YY1 also presented puncta overlapping with endogenous proteins of the three coactivators (Figure 5B), but with significantly increased sizes versus those of endogenous YY1 (Figure 5C). Ser2 and Ser5 phosphorylation of the C-terminal heptapeptide repeats of RNA polymerase II (Pol II S2P and S5P) are general markers of gene transcription (42). Endogenous YY1 colocalized with Pol II S2P and S5P signals (Figure 5D), suggesting that YY1 promotes gene transcription. Similar to coactivators, Flag-YY1 showed relatively intensified puncta overlapping with Pol II S2P and S5P signals (Figure 5E) with increased sizes versus those by endogenous YY1 (Figure 5F). In line with these data, mCherry- YY1 could form overlapped droplets with EGFP-fused IDR regions of MED1, BRD4 and Pol II *in vitro* (Figure 5G). Based on these results, we propose that YY1 plays a key role in promoting gene transcription through recruiting major coactivators to form phase separation. Consistent with this hypothesis, both endogenous YY1 and Flag-YY1 showed overlapped puncta with gene activation histone markers H3K27ac, H3K4me1 and H3K4me3, but not a repressive marker H3K9me3 (Figures 5H and 5I). In addition, Flag YY1 markedly increased sizes of the puncta overlapping with the three activation markers but not H3K9me3 (Figure 5J). Immunostaining assays were also conducted in MCF-7 cells with virtually the same results (Supplementary Figures S4A-S4I). Overall, our data strongly suggested that YY1 is a general transcriptional activator to promote gene expression.

**Figure 5.**
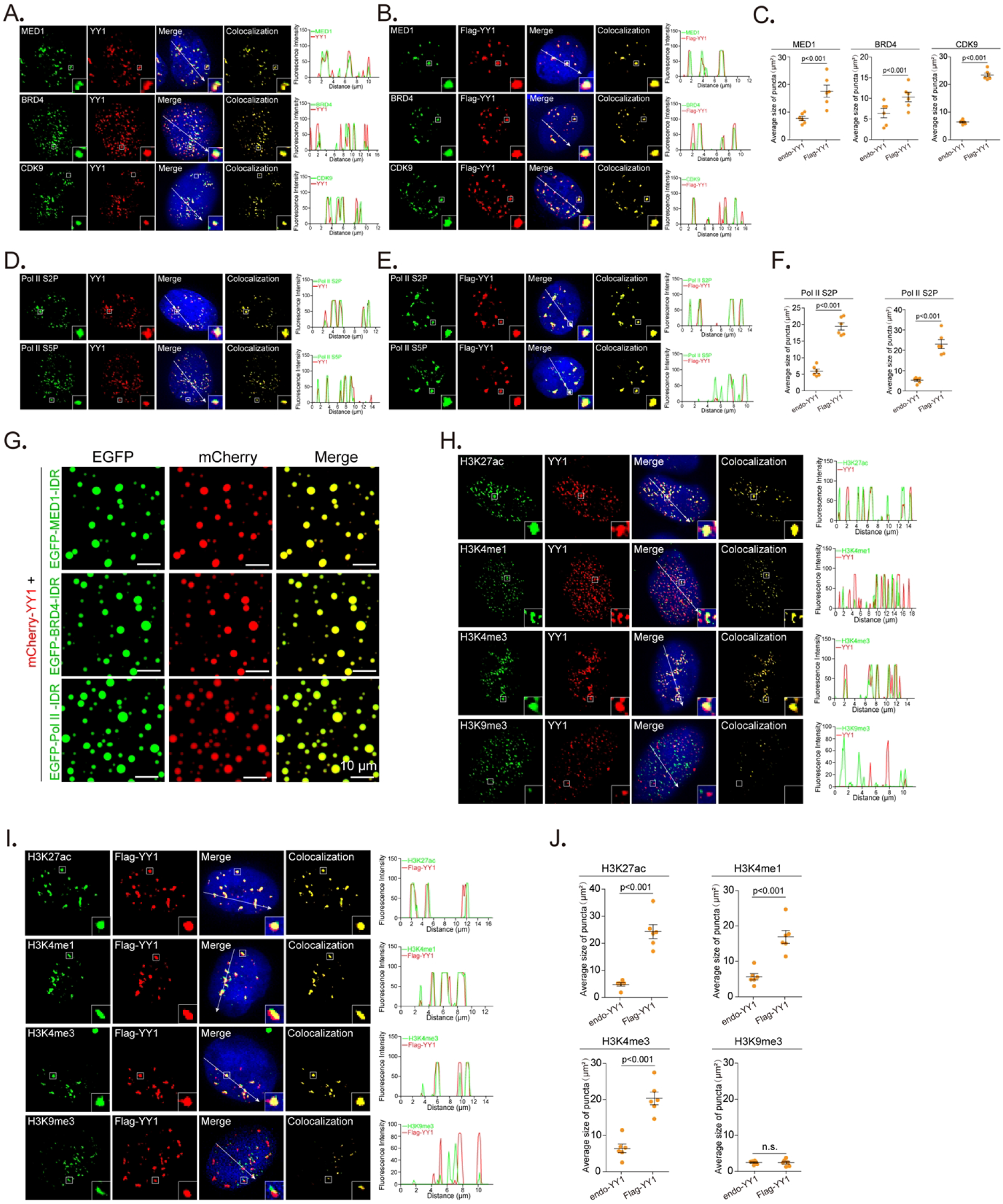
YY1 compartmentalizes additional coactivators to nuclear puncta. **A** and **B**. Colocalization of BRD4, MED1 and CDK9 with endogenous YY1 (**A**) or with Flag-YY1(**B**) in nuclear puncta in MDA-MB-231 cells. Endogenous proteins were detected by their corresponding antibodies. Flag-YY1 was detected by a Flag epitope antibody. Nuclei were visualized by DAPI staining. Line scans of the colocalization images are depicted by white arrows with quantification shown at right. **C**. Quantified average sizes of merged puncta in MDA-MB-231 cells with endogenous YY1 (A) and Flag-YY1 (B). Data are mean ± s.e.m. of puncta in 6 fields in each group. **D** and **E**. Colocalization of active RNA Pol II with YY1 (**D**) or Flag-YY1 (**E**) in nuclear puncta of MDA-MB-231 cells. Active RNA Pol II was detected by antibodies for phosphorylation of Ser 5 (S5P) or Ser 2 (S2P), with nuclei detected by DAPI. Line scans of colocalization images are depicted by white arrows with quantification shown at right. **F**. Quantification of average sizes of merged puncta in MDA-MB-231 cells with endogenous YY1 (D) and Flag-YY1 (E). Data are presented as mean ± s.e.m. from puncta of 6 fields in each group. **G**. Representative images of droplet formation of mCherry-YY1 with EGFP- MED1-IDR, EGFP-BRD4-IDR, or EGFP-Pol II-IDR. **H** and **I**. Localization of active (H3K27ac, H3K4me1 and H3K4me3) and repressive (H3K9me3) histone markers with endogenous YY1 (**H**) or Flag-YY1 (**I**) in MDA-MB-231 cells. Histone markers were determined using corresponding antibodies. Nuclei were detected by DAPI. Line scans of colocalization images are depicted by white arrows with quantification shown at right. **J**. Quantified average sizes of merged puncta in MDA-MB-231 cells with endogenous YY1 (H) and Flag-YY1 (I). Data are presented as mean ± s.e.m. of puncta in 6 fields in each group. All experiments in this figure were independently repeated at least 6 times with similar results.

### YY1 activates FOXM1 gene expression through recruiting general coactivators

YY1 conditional knockout in mouse embryonic fibroblast cells revealed many genes as potential targets of YY1 (43). Among them, FOXM1 exhibited over 2-fold reduction in response to YY1 depletion. Both YY1 and FOXM1 play oncogenic or proliferative roles in tumorigenesis (16,44). Consistently, they were overexpressed in breast cancer and associated with patients’ poor prognosis (38,45). Analyses of a TCGA dataset of mammary samples indicated positive YY1 and FOXM1 correlations (Supplementary Figure S5A), especially in normal tissues, suggesting physiological significance of their functional interplay. In mammary cells, we detected concurrent increase of YY1 and FOXM1 expression in breast cancer cells versus nontumorigenic MCF-10A cells (Figure 6A). At both mRNA and protein levels, ectopically expressed YY1 in MCF-10A cells increased endogenous FOXM1 expression, while shRNA-mediated YY1 knockdown reduced it in breast cancer cells (Figure 6B). All these data strongly suggest a positive regulation of FOXM1 gene by YY1.

**Figure 6.**
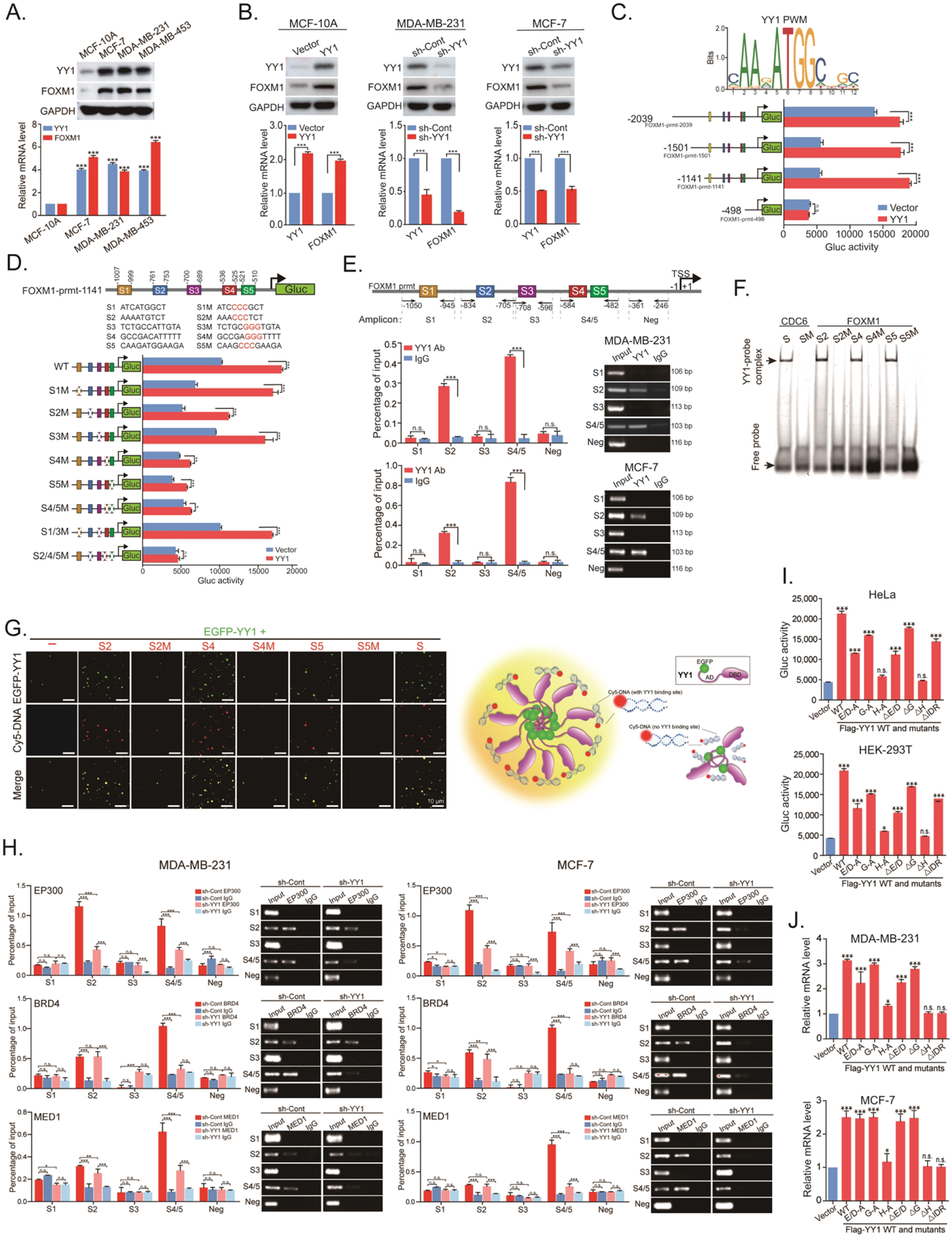
YY1 binds to the FOXM1 promoter and activates its expression. **A**. YY1 and FOXM1 expression in mammary cell lines. Nontumorigenic MCF-10A cells, and three indicated breast cancer cell lines were analyzed by Western blot using indicated antibodies (top row). YY1 and FOXM1 mRNA levels were quantified by RT-qPCR (bottom row). **B**. Effects of YY1 on FOXM1 in mammary cells. MCF-10A cells were infected by lentivirus expressing YY1, while MDA-MB-231 and MCF-7 cells were infected by lentivirus carrying shYY1. YY1 and FOXM1 protein and mRNA levels were analyzed by Western blot (top row) and RT-qPCR (bottom row). **C**. Mapping YY1- regulated region in the FOXM1 promoter. Conserved YY1 binding elements with a core sequence of ATGG (Jaspar Matrix ID: MA0095.2, top row). Reporters with different FOXM1 promoter lengths driving Gluc were generated. Data of reporter assays in response to ectopic YY1 in HeLa cells were examined (bottom row). Data represent mean ± s.d. (n = 3). **D**. Evaluation of putative YY1 binding sites in the FOXM1 promoter. Putative YY1 binding sites S1-S5 were identified in the 1141-bps region upstream of the translational start site (TSS) of the FOXM1 gene, with WT and mutants presented (top panel). Data of reporter assays in response to YY1 in HeLa cells are presented in bottom row. **E**. ChIP-qPCR assays to examine YY1 binding to the FOXM1 promoter. The FOXM1 promoter is presented in top row with five qPCR primer pairs. YY1 antibody and a control IgG were used in ChIP assay. Data quantification and agarose gels of ChIP-qPCR in MDA-MB-231 and MCF-7 cells were presented in middle and bottom rows, respectively. **F**. EMSA to test YY1 binding to the FOXM1 promoter. His×6-YY1 was incubated with Cy5-labeled S2, S4, S5 probes, and mutants (S2M, S4M and S5M) with a CDC6 promoter probe and its mutant as positive and negative controls, respectively. YY1-probe complex and free probe positions are label on the left. **G**. Left: representative images of droplet formation by EGFP-YY1 with Cy5-labeled probes in Figure 6F and Supplementary Table S2. Right: schematic model of droplet formation promoted by DNA probes. **H**. ChIP-qPCR assays to test coactivators’ binding to the FOXM1 promoter. EP300, BRD4 and MED1 antibodies, and a control IgG were used in ChIP assays, and qPCR primers are shown in Supplementary Table S1. Both quantification and agarose gels of ChIP-qPCR in MDA-MB-231 and MCF-7 cells were presented. **I** and **J**. Evaluation of YY1 mutations’ effects on the FOXM1 promoter. In **I**, FOXM1 promoter reporter (pFOXM1-prmt-Gluc) and pCMV-SEAP vector were cotransfected with YY1 WT and mutant vectors into HeLa (top) and HEK-293T (bottom) cells in 24-well plates. Gluc activity in each well was measured and normalized against its SEAP activity. In **J**, YY1 vectors were transfected into MDA-MB- 231 (top) and MCF-7 (bottom) cells, followed by RT-qPCR to quantify FOXM1 mRNA levels.

To examine mechanism regulating FOXM1 gene expression, we first mapped essential region of the FOXM1 promoter. Four reporter constructs were generated with Gaussia luciferase (Gluc) driven by different lengths of the upstream sequence from the TSS of the FOXM1 gene (Figure 6C). Based on response of the reporters to cotransfected YY1, YY1-regulated essential elements reside within the 1,141-bps region upstream of the TSS (Figure 6C). To explore potential regulation of FOXM1 expression by YY1, we examined the human FOXM1 promoter for YY1 consensus binding elements using the JASPAR (46) and Tfsitescan (http://www.ifti.org/cgi-bin/ifti/Tfsitescan.pl) databases. Within the 1,141-bps FOXM1 promoter, we identified five potential YY1 binding sites (Supplementary Figure S5B), and mutagenesis of #2, #4 and #5 sites, especially the latter two, caused remarkable reduction of FOXM1 promoter activity in reporter assays (Figure 6D). YY1 binding to the FOXM1 promoter through these sites was verified by chromatin immunoprecipitation (ChIP) assay (Figure 6E). To test YY1 binding *in vitro*, we synthesized double-stranded (ds) oligonucleotides S1 to S5 based on corresponding YY1 binding sites in the FOXM1 promoter. In EMSA studies, a Cy5-labeled probe based on a YY1 consensus motif in the CDC6 promoter (47) could be out-competed in His×6-YY1 binding by S2, S4 and S5, but not S1 and S3, or mutants of S2, S4 and S5 (Supplementary Figure S5C). In addition, Cy5-labeled S2, S4 and S5, but not their mutants, could bind YY1 to form slowly migrated bands in EMSA (Figure 6F). To evaluate the potential of YY1-regulated FOXM1 expression through a phase separation mechanism, we incubated Cy5-labeled oligonucleotides with a relatively low concentration (2 µM) of EGFP-YY1. Strikingly, the presence of Cy5-labeled S2, S4 and S5, but not their mutants, could associate with EGFP-YY1 to form droplets with relatively large sizes (Figure 6G, left panel). The data not only suggested a phase separation mechanism regulating FOXM1 gene transcription, but also disclosed an important fact that YY1 phase separation can be promoted by binding to DNA containing its consensus motifs (Figure 6G, right panel). To evaluate whether super-enhancer was involved in regulating FOXM1 expression, we carried out ChIP assays for several coactivators. Both semi-quantitative and quantitative PCR analyses revealed that EP300, BRD4 and MED1 could bind to the regions containing YY1’s S2 and S4/5 consensus sites, but not those of S1 and S3 sites, in the FOXM1 promoter in breast cancer cells (Figure 6H). Consistent with the observation above, both YY1-H-A and -ΔH mutants, deficient in phase separation, exhibited the least capability in promoting both promoter reporter and endogenous transcript of the FOXM1 gene (Figures 6I and 6J). Interestingly, in reporter assays of Figure 6I, transfection of YY1-ΔIDR still retained significant Gluc activity, but its expression in two breast cancers could not activate the FOXM1 gene (Figure 6J). Noteworthily, endogenous YY1 was still present in cells of reporter assays, but knocked down when testing YY1 mutants in breast cancer cells. Thus, endogenous YY1 was likely involved in driving reporter expression, which could be functionally perturbed by YY1 mutants with deficient IDRs, but not the mutant with ΔIDR. Nevertheless, ectopic YY1-ΔIDR could still occupy the promoters of many reporters and thus reduced overall Gluc expression (Figure 6I).

### YY1 promotes the formation of super-enhancer to drive FOXM1 gene expression

YY1 has been reported to regulate gene expression through promoting enhancer activity (27,48,49), while phase separation is a characteristic feature of enhancer mechanism (50). To interrogate whether enhancers were involved in YY1-regulated FOXM1 expression, we surveyed the regions of overlapped YY1 binding enrichment and gene activation markers in the vicinity of the FOXM1 gene in the human genome. Within 650 kbs of the FOXM1 promoter, we identified five candidate enhancer regions, and designated them as E1 to E5 (Figure 7A). The chromosome conformation capture (3C) approach (32) with restriction enzyme digestion, digested genomic DNA ligation, and PCR amplification of ligated DNA, was used to examine vicinal regions in forming enhancer complexes through interacting with the FOXM1 promoter. Multiple EcoRI-digested segments displayed ligation-dependent PCR bands (Figure 7B), mostly overlapping with enhancers E3, E4 and E5 (Figure 7A). Similarly, the genomic DNA digested by HindIII also showed positive PCR bands overlapping with E3, E4 and E5, depending on T4 DNA ligase (Supplementary Figure S6A). Furthermore, shRNA-mediated YY1 knockdown eliminated these PCR bands (Supplementary Figure S6B), indicating that physical closeness of the enhancer elements to the FOXM1 promoter was contingent to the presence of YY1. Importantly, we confirmed the precise ligation between digested segments of the FOXM1 promoter and the enhancers by DNA sequencing analysis (Supplementary Figure S6C). In addition, 1,6-hexanediol treatment greatly reduced PCR products (Figure 7C), suggesting that phase separation is prerequisite for FOXM1 promoter’s proximity to enhancers. To assess potential enhancer properties of EcoRI-digested segments 19, 20, 26, 27 and 29, we generated FOXM1 promoter reporters with sub-segmented sequences of these regions located either upstream or downstream according to their natural positions relative to the FOXM1 TSS in the genome (Figure 7D, top row). In reporter assays, several fragments from these segments showed response to YY1’s ectopic expression or knockdown (Figure 7D, bottom row), consistent with their enhancer identity. In line with these data, treatment of transfected cells by JQ1, an enhancer inhibitor, dampened reporter activities (Figure 7E).

**Figure 7.**
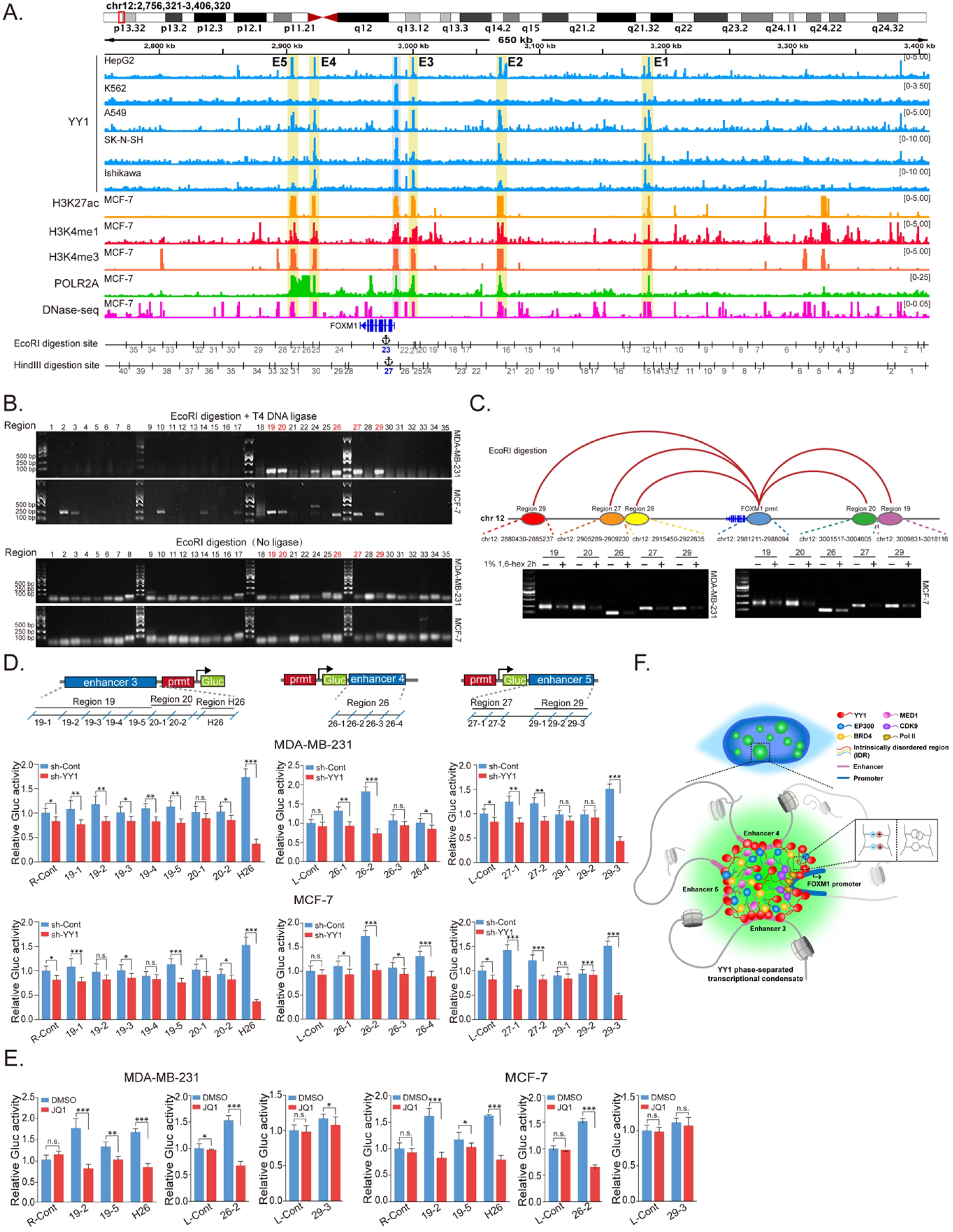
YY1 phase-separated transcription complex connects distal enhancer elements and the FOXM1 promoter. **A.** Schematic view across the FOXM1 gene locus (chr12:2,768,321-3,408,320 [hg19]) with genomic and epigenetic information. Graphic active regulatory regions were generated using the ENCODE database, and potential enhancers (E1 to E5) are shaded in yellow. Fragments digested by EcoRI or HindIII are numbered and shown below the graph. **B**. Chromatin conformation capture (3C) analysis to examine direct physical interactions between the FOXM1 promoter and enhancer elements. The 3C analyses followed the protocol in Materials and Methods, using EcoRI in genomic DNA digestion. Ligation step was performed with or without T4 DNA ligase as indicated. Numbers on gel top correspond to “EcoRI digestion site” in Figure 8A. PCR to examine direct physical interactions between the FOXM1 promoter and each of EcoRI fragments used the primer sets in Supplementary Table S3. Numbers of fragments overlapping with enhancers and interacting with the FOXM1 promoter are in red text. **C**. Schematic interactions between enhancers and the FOXM1 promoter on chromosome 12 (top row), and PCR product images of ligated DNA in 3C assays conducted in the absence or presence of 1% of 1,6-hexanediol (bottom row). **D**. Reporter assays to examine the effects of enhancers on the FOXM1 promoter. Sectionalized enhancers 3, 4 or 5 were individually placed adjacent to the FOXM1 promoter to create reporter vectors, as schematically shown in top row. Reporter vectors were transfected into MDA-MB-231 and MCF-7 cells with or without YY1 knockdown (bottom row). Data are shown as mean ± s.d. (n=3). **E**. Reporter assays to assess the effects of JQ1. Reporter vectors with enhancer fragments were transfected into MDA-MB-231 and MCF-7 cells with or without JQ1, followed by examining Gluc activity. Data are shown as mean ± s.d. (n=3). **F**. Schematic model of a super-enhancer formed by YY1-mediated phase separation with incorporation of coactivators and stabilization by three distal enhancers to activate FOXM1 gene transcription.

We further evaluated the importance of YY1-regulated FOXM1 expression in mammary cells. As we previously reported, ectopic YY1 promoted proliferation of primary mammary epithelial cells, while its depletion reduced it (38). In the current study, FOXM1 knockdown or its ectopic expression could significantly counteract cell viability changes in MCF-10A cells (by exogenous YY1) or breast cancer cells (by YY1 silencing), respectively (Supplementary Figure S7A). We reported YY1-promoted AKT activation (51); consistently, FOXM1 could reinstate AKT-T308 and -S473 phosphorylation reduced by YY1 knockdown (Supplementary Figure S7B). FOXM1 depletion attenuated YY1-promoted migration and clonogenicity of MCF-10A cells in wound healing and colony formation assays, while, in these assays, ectopic FOXM1 could significantly rescue the deficiencies of breast cancer cells caused by YY1 silencing (Supplementary Figures S7C and S7D).

Based on our data, we propose a model of YY1-regulated gene activation (Figure 7F). In this model, a super-enhancer is formed by YY1-mediated phase separation that comprises major coactivators and is stabilized by three distal enhancers to activate FOXM1 promoter and stimulate its gene transcription.

## DISCUSSION

Phase separation is a general phenomenon in polymer chemistry, but has recently been developed into a concept or mechanism of biological regulation (52). Liquid-liquid phase separation in different subcellular sections creates membrane-less condensates that compartmentalize biomolecules, such as proteins, RNAs and DNAs, with pertinent biological activities, and allows a specific biological event to be processed in a relatively undisturbed manner (13). Applications of phase separation in transcriptional regulation are the seminal discovery of Young’s group through demonstrating the formation of phase-separated condensates that confine various coactivators and function as super-enhancers (8,9). These discoveries largely extended our view of transcriptional regulation and revolutionized mechanism or concept of sustained gene expression through super-enhancers.

As a key transcription factor, YY1 interacts with numerous epigenetic writers and erasers (16). In addition to recognizing its consensus sites, YY1 binds G-quadruplex structures (53). Importantly, YY1’s ability in homodimerization (27,53) allows it to bridge promoters and enhancer elements. Weintraub *et al*. reported that YY1 generally occupies active enhancers and promoters in different cell types, and perturbation of YY1 binding disrupts enhancer-promoter looping (27). The promiscuous interaction of YY1 with coactivators can promote their recruitment into the enhancer complexes. Consistently, a number of studies demonstrated YY1’s participation in forming enhancers or super-enhancers to regulate genes involved in various biological processes (27,49,54-56). These studies strongly support a key role of YY1 in promoting super-enhancer assembly to activate gene expression. However, despite these exciting indications or hints to us, molecular evidence is still lacking for detailed mechanisms of YY1-regulated enhancer formation.

YY1 protein primary structure has several unique features scarcely observed in other proteins, including a highly acidic N-terminus, consecutive E/D-, G- and H-rich regions in the TAD, and four tandem zinc fingers at a basic C-terminus, as well as a high lysine content (32 versus total 414 amino acids) but none of them found in first 157 residues. When scanning YY1 sequence, we discover high propensity of structural disorder in its TAD. Subsequently, our experimental data unequivocally demonstrate YY1’s competence in undertaking liquid-liquid phase separation both *in vitro* and in cell. Interestingly, we identify the 11×H cluster as a critical motif of YY1 to promote its phase separation, which can be extended to four additional histidine cluster-containing proteins.

The YY1-rich nuclear phase-separated condensates are likely super-enhancers based on their participation by major coactivators, including EP300, MED1, BRD4 and CDK9, as well as active Pol II. With FOXM1 expression as a regulatory model, we also demonstrate the role of YY1 in convening three distal enhancer elements and the FOXM1 promoter to assemble a super-enhancer for gene activation. Strikingly, oligonucleotides containing YY1 consensus sites in the FOXM1 promoter can participate in and steadily facilitate YY1 droplet formation *in vitro*, suggesting that YY1-mediated phase-separated nuclear condensates are likely involved and promoted by genomic DNA. Interestingly, Sigova, *et al*. demonstrated that YY1 could bind both gene regulatory elements and their associated RNA species transcribed from enhancers and promoters in a genome-wide manner (17). Thus, it is logical to predict that cognate RNAs may participate YY1-coordinated phase separation, which deserves future investigation.

Despite mounting studies showing that YY1 either activates or represses gene expression, we observe YY1 colocalization with coactivators and histone markers for gene activation, but not for repression. Our data are consistent with previous studies showing YY1’s general association with active promoters and enhancers (17,27). Thus, although YY1 was reported to suppress gene expression through recruiting corepressors, such as HDACs, EZH2 and DNMTs (57-59), our data support its primary role as a transcriptional activator.

FOXM1 is recognized as a critical proliferation-associated transcription factor, regulating cell proliferation, self-renewal, and tumorigenesis. As a proliferative gene, FOXM1 overexpression was reported in almost all cancer types, and correlated with patients’ poor prognoses (44). Consistently, FOXM1 gene is regulated by a variety of oncogenic TFs, such as E2Fs, MYC, ERα, STAT3 and CREB, as well as FOXM1 itself (44). In the current study, we provide the first evidence to demonstrate FOXM1 expression promoted by super-enhancers formed by YY1-containing phase-separated condensates, which extends molecular mechanisms driving FOXM1 overexpression in cancers. Interestingly, FOXM1 has also been reported to participate enhancer regulation of other genes (60,61). Based on its self-regulated feature, FOXM1 itself may potentially get involved in super-enhancer regulating its expression.

Our provide both *in vitro* and cell-based data to demonstrate capability of EP300 in forming phase separation. Based on algorithm prediction, we identify five potential IDRs in EP300 primary sequence, and verify that IDR3 (at middle region) and IDR5 (at C-terminus) form droplets *in vitro*. In two recent studies, EP300 C-terminus was reported to undertake phase separation (62,63), consistent with our results, but Ma, *et al*. showed the capability of its N-terminus (1-566 amino acids) in forming droplets *in vitro*, which was not the case in our study. Nevertheless, we provided ample evidence to demonstrate the competence of EP300 in forming phase-separated condensates, including the optimal parameters, and proved its role as a primary coactivator of YY1-mediated enhancer complex.

## SUPPLEMENTARY DATA

Supplementary Data are available at NAR online.

## ACKNOWLEDGEMENT

WW, DL and GS initiated the study, designed the experiments, analyzed data, and wrote the manuscript; WW, SQ, CY, CZ and DL carried out the research; GL also analyzed data; DBS and JS contributed conceptual suggestions and new reagents/analytic tools.

