## Supplemental File for "A histidine cluster determines YY1-compartmentalized coactivators and chromatin elements in phase-separated super-enhancers"

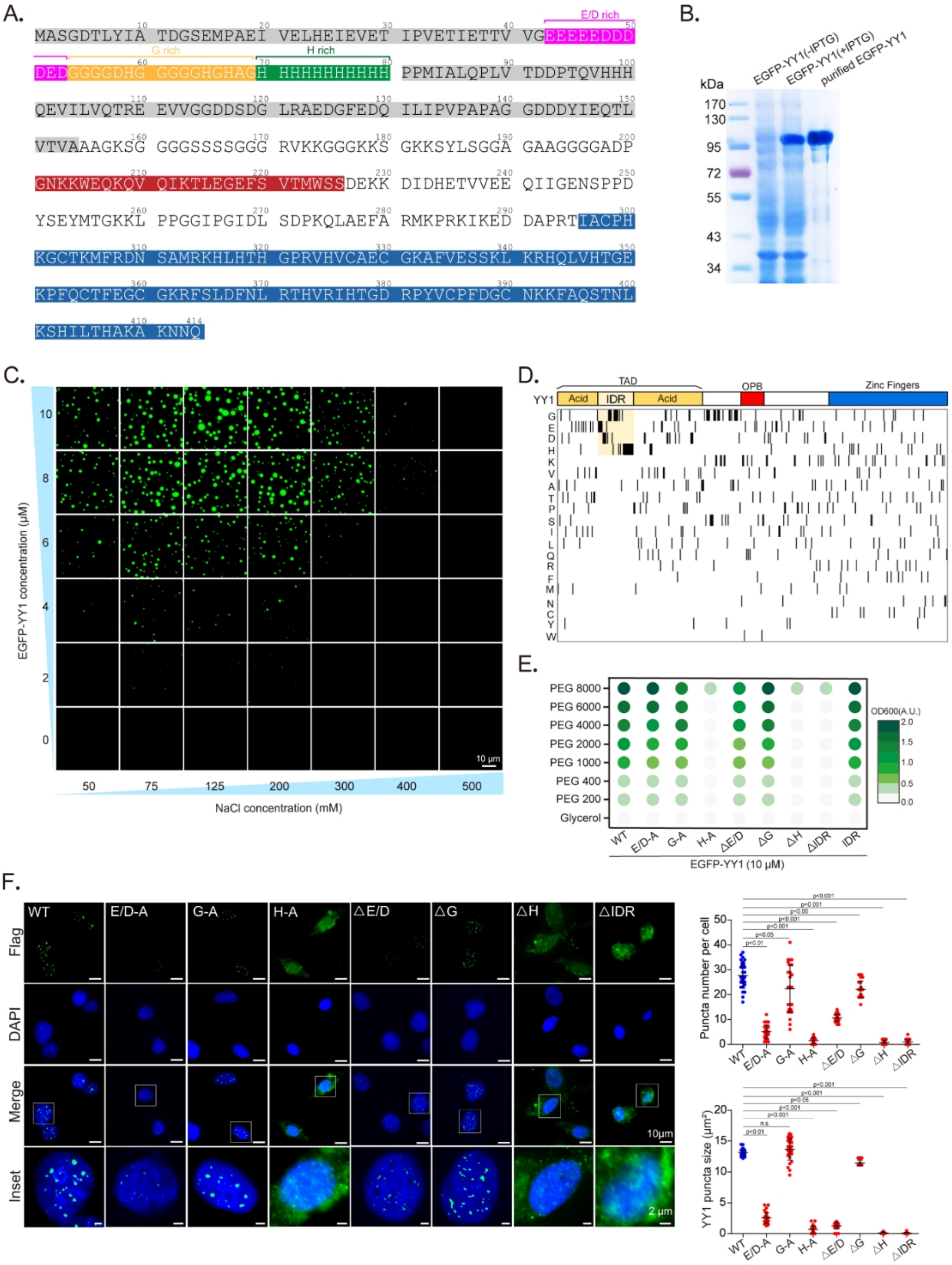

**Figure S1: Sequence characteristics and phase separation conditions of YY1 protein.** (**A**) YY1 primary sequence and its specific features. The glutamic and aspartic acid rich region is in magenta color, glycine rich region is in yellow color, histidine rich region is in green color, acid region is in grey color, OPB region is in red color, and zinc finger sequence is in blue color. The YY1 sequence was retrieved from the UniProt database (entry ID: P25490). (**B**) SDS-PAGE of lysate supernatants from bacteria expressing His×6-EGFP-YY1 vector without or with isopropyl β-D-1-thiogalactopyranoside (IPTG) induction, and purified His×6-EGFP-YY1. (**C**) Representative images of EGFP-YY1 droplet formation at different protein and NaCl concentrations. Assays were repeated in six independent experiments. (**D**) Enrichment analysis of YY1 protein's amino acid in an order of frequencies. Each row represents position of a single amino acid. (**E**) Concentration-dependent increase of turbidity in the solutions of EGFP-YY1 WT or mutants (10 μM each) with increasing molecular weights of PEGs. (**F**) Representative nuclear images of U2OS cells transfected by 3×Flag-YY1 WT and mutant vectors. Flag-YY1 was detected by immunofluorescence staining using a Flag antibody. Nuclei were detected by DAPI staining (left panel). Thirty cells were quantified for numbers and sizes of puncta using Image J. Data represent mean ± s.e.m. (right panel).

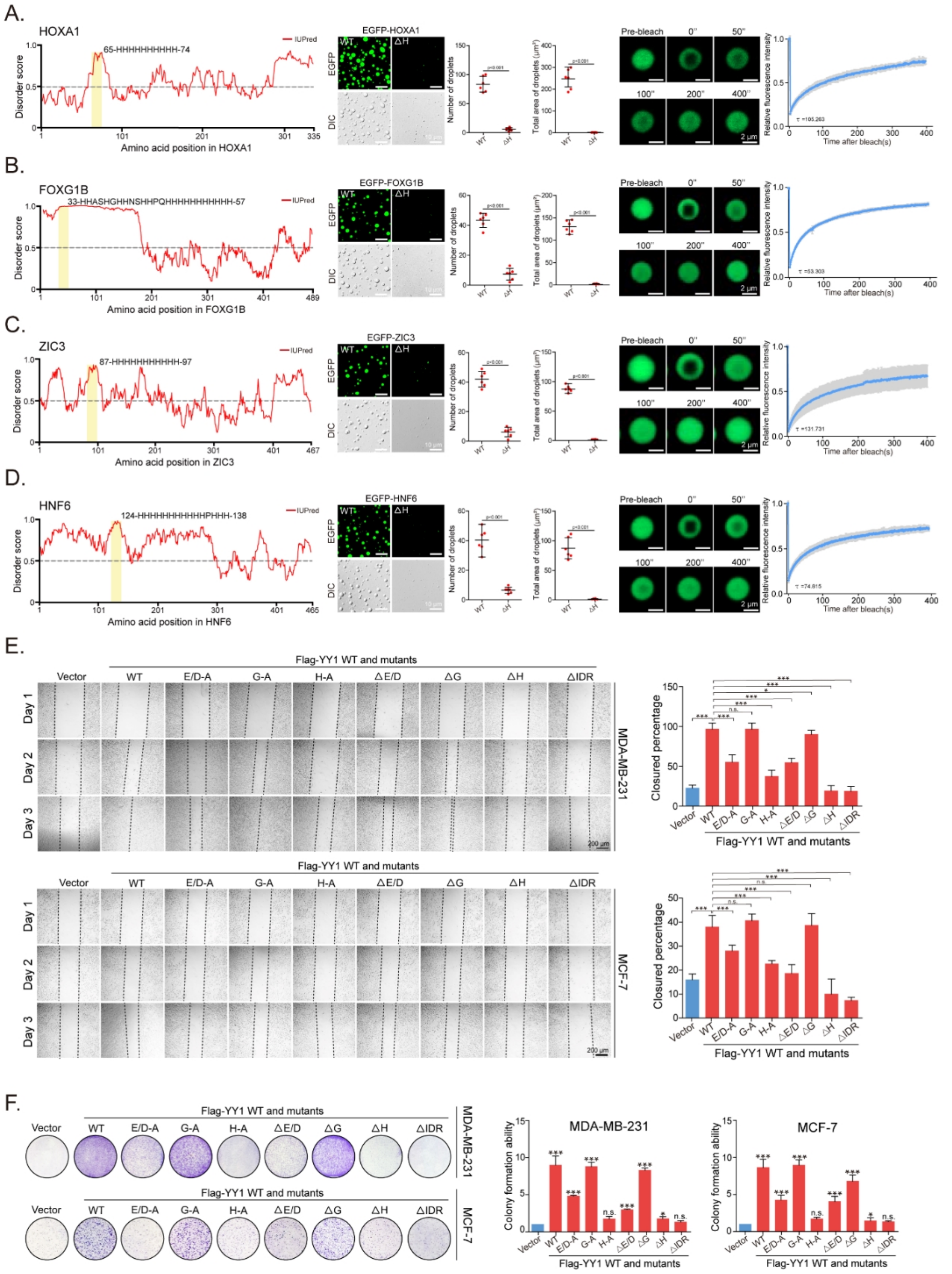


**Figure S2： Contributions of His-cluster to LLPS of four addition proteins and the effects of YY1 TAD mutations on its function.** (**A** to **D**) Graphs of intrinsic disorder of histidine cluster-containing proteins HOXA1 (A), FOXG1B (B), ZIC3 (C) and HNF6 (D), as calculated by the IUPred (http://iupred.elte.hu/) algorithm. The scores are assigned between 0 and 1, and a score above 0.5 indicates disorder. Yellow shades indicate histidine-rich domains (left panels). Representative images of droplet formation of each EGFP-fusion protein (4 μM) in the droplet formation buffer containing 125 mM NaCl and 10% PEG-8000, and their quantification (mean ±

s.e.m., n = 6) are shown in middle panels. FRAP recovery images and curves of the droplets formed by the four purified recombinant EGFP-fusion proteins are presented at the right panels. (**E**) and (**F**) The same treatments of MDA-MB-231 and MCF-7 cells with Figures 2G and 2H to test the effects of YY1 mutants on cell migration in wound healing assays (E) and clonogenicity in colony formation assays (F). Quantification is shown at the right panels.

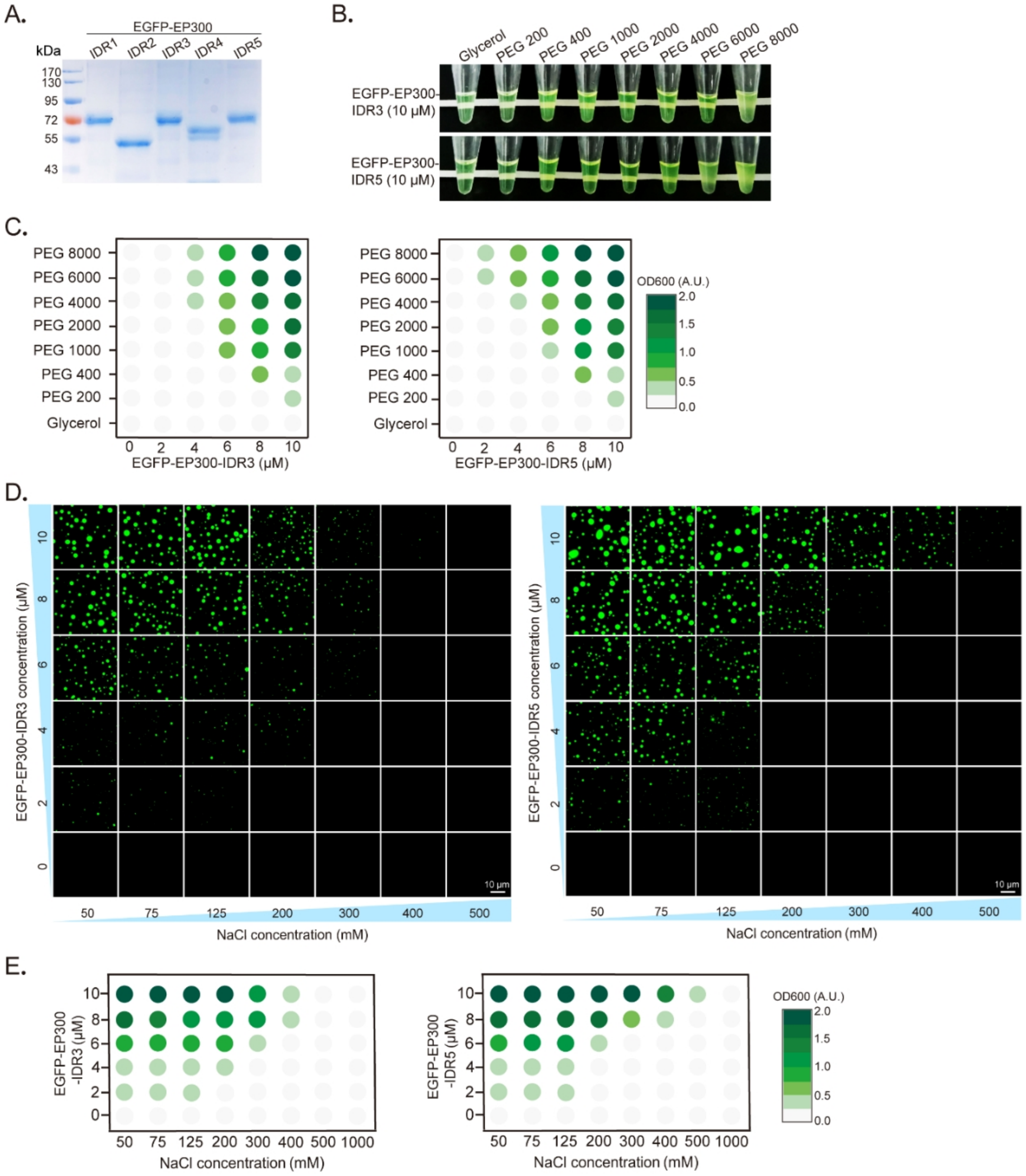


**Figure S3: Additional characterizations of EP300's phase separation capability in vitro.** (**A**) SDS-PAGE analysis of purified recombinant EGFP-EP300-IDR proteins expressed in E. coli. (**B**) Visualization of turbidity associated with droplet formation. EP300-IDR3 and EGFP-EP300-IDR5 (10 μM of each) were individually incubated in the droplet formation buffer containing 125 mM NaCl and PEGs with increasing molecular weights to mimic large polymeric crowders. (**C**) Phase diagram of OD600nm showing turbidity changes caused by EGFP-EP300-IDR3 (left panel) and EGFP-EP300-IDR5 (right panel) droplet formation in PEGs with increasing molecular weights and protein concentrations. (**D**) Representative images showing droplet formation of EGFP-EP300-IDR3 (left panel) and EGFP-EP300-IDR5 (right panel) at different protein and NaCl concentrations. Data are representative of six independent experiments. (**E**) Phase diagram of OD600nm showing turbidity changes caused by EGFP-EP300-IDR3 (left panel) and EGFP-EP300-IDR5 (right panel) droplet formation at different protein and NaCl concentrations.

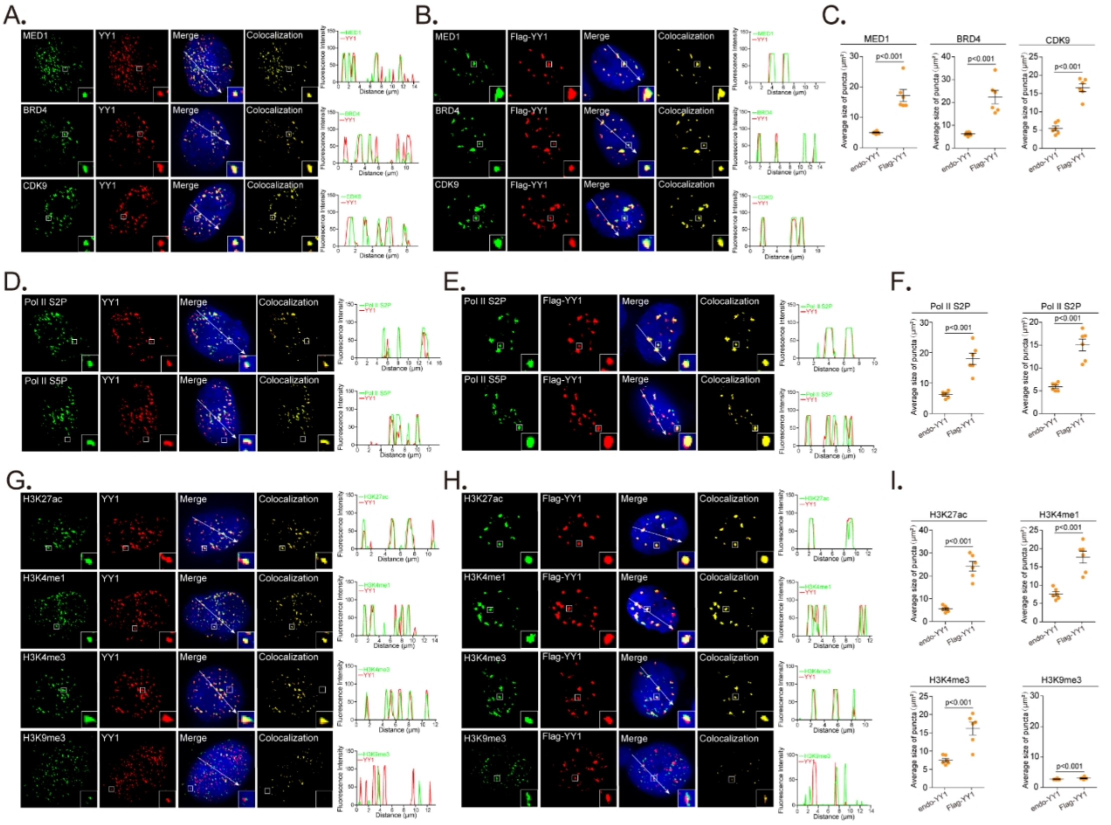


**Figure S4： YY1 compartmentalizes major transcription coactivators to nuclear puncta in MCF-7 cells.** (**A** and **B**). Colocalization of endogenous BRD4, MED1 and CDK9 with endogenous YY1 (A) or with ectopic Flag-YY1(B) in nuclear puncta of MCF-7 cells. Localization of endogenous MED1, BRD4, CDK9 and YY1 was detected by immunofluorescence staining using their corresponding antibodies. Flag-YY1 was detected by a Flag antibody and nuclei were visualized by DAPI. Line scans of the images of BRD4, MED1 or CDK9 and YY1 or Flag-YY1 colocalization at the positions are depicted by white arrows with quantification shown at right panel. (**C**) Quantification of average sizes of merged puncta in MCF-7 cells with endogenous YY1 (A) and Flag-YY1 (B). Data are presented as mean ± s.e.m. from puncta in six fields of each group. (**D** and **E**). Colocalization of active RNA Pol II with endogenous YY1 (D) or Flag-YY1 (E) in nuclear puncta of MCF-7 cells. Localization of active RNA Pol II was detected by immunofluorescence staining using antibodies against phosphorylation of Ser 5 (S5P) or Ser 2 (S2P). Endogenous YY1 and Flag-YY1 was detected by immunofluorescence staining using antibodies against YY1 and Flag epitope, respectively. Nuclei were detected by DAPI. Line scans of the colocalization images at the positions are depicted by white arrows with quantification shown at right panel. (**F**) Quantification of average sizes of merged puncta in MCF-7 cells with endogenous YY1 (D) and Flag-YY1 (E). Data are presented as mean ± s.e.m. from puncta in six fields of each group. (**G** and **H**) Relative localization of active (H3K27ac, H3K4me1 and H3K4me3) and repressive (H3K9me3) histone markers with endogenous YY1 (G) or Flag-YY1 (H) in the nuclear puncta of MCF-7 cells. Levels of histone markers were tested by immunofluorescence staining using corresponding antibodies. Nuclei were detected by DAPI. Line scans of the colocalization images at the positions are depicted by white arrows with quantification shown at right panel. (**I**) Quantification of average sizes of merged puncta in MCF-7 cells with endogenous YY1 (G) and Flag-YY1 (H). Data are presented as mean ± s.e.m. of puncta in six fields in each group. All experiments in this figure were independently repeated at least six times with similar results.

A.

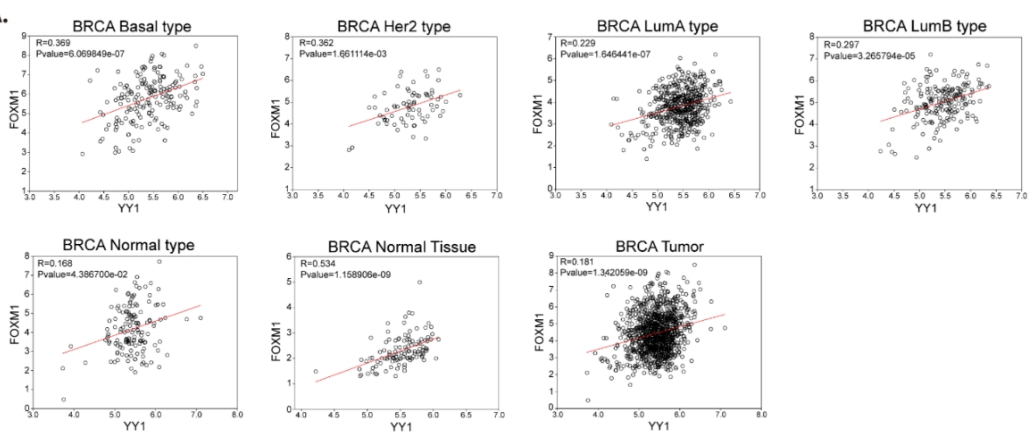


B.

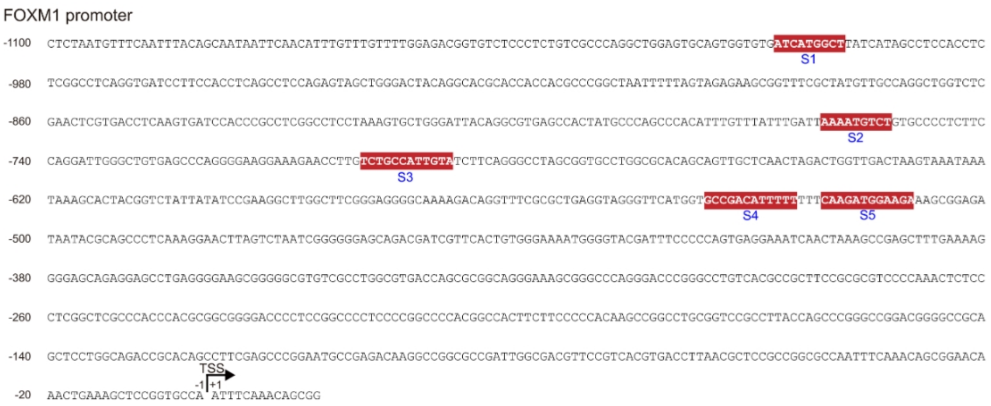


| # | YY1 binding sites | positions of the core sites | websites |
|---|---|---|---|
| S1 | ATCATGGCT | -1007 to -999 | Tfsitescan (http://www.ifti.org/cgi-bin/ifti/Tfsitescan.pl) |
| S2 | AAAATGTCT | -761 to -753 | Tfsitescan (http://www.ifti.org/cgi-bin/ifti/Tfsitescan.pl) |
| S3 | TCTGCCATTGTA | -700 to -689 | JASPAR (http://jaspar.genereg.net/) |
| S4 | GCCGACATTTTT | -536 to -525 | Tfsitescan (http://www.ifti.org/cgi-bin/ifti/Tfsitescan.pl) & JASPAR (http://jaspar.genereg.net/) |
| S5 | CAAGATGGAAGA | -521 to -510 | JASPAR (http://jaspar.genereg.net/) |

C.

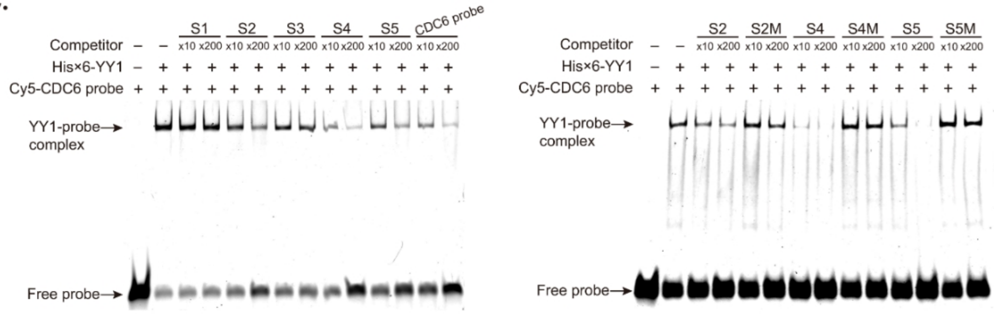


**Figure S5: Positive YY1 and FOXM1 correlation in mammary tissues and YY1 binding site analyses in the FOXM1 promoter.** (**A**) Correlative analyses between YY1 and FOXM1 expression from a TCGA-BRCA dataset (Cancer Genome Atlas Network. ***Nature***. 490:61-70) using the breast cancer subtypes categorized by the PAM50 gene signature. (**B**) DNA sequence from -1100 to -1 in the FOXM1 promoter with five putative YY1 binding sites (S1-S5) highlighted in a red background. The transcription start site (TSS) of the FOXM1 gene is indicated, and the arrow shows the direction of transcription (top panel). Predicted YY1 binding sites in the FOXM1 promoter by the Tfsitescan or JASPAR website are displayed at the bottom panel. (**C**) EMSA to test competitive binding of FOXM1 promoter oligonucleotides and a labeled CDC6 promoter probe to YY1 protein. His×6-YY1 (1 μg) and 0.5 pmol of Cy5-labeled CDC6 probe were mixed, with addition of 10- or 200-fold of unlabeled WT or mutant probes for competitive binding, as indicated on the top. The sequences of S1-S5, S2M, S4M and S5M are shown in table S2.

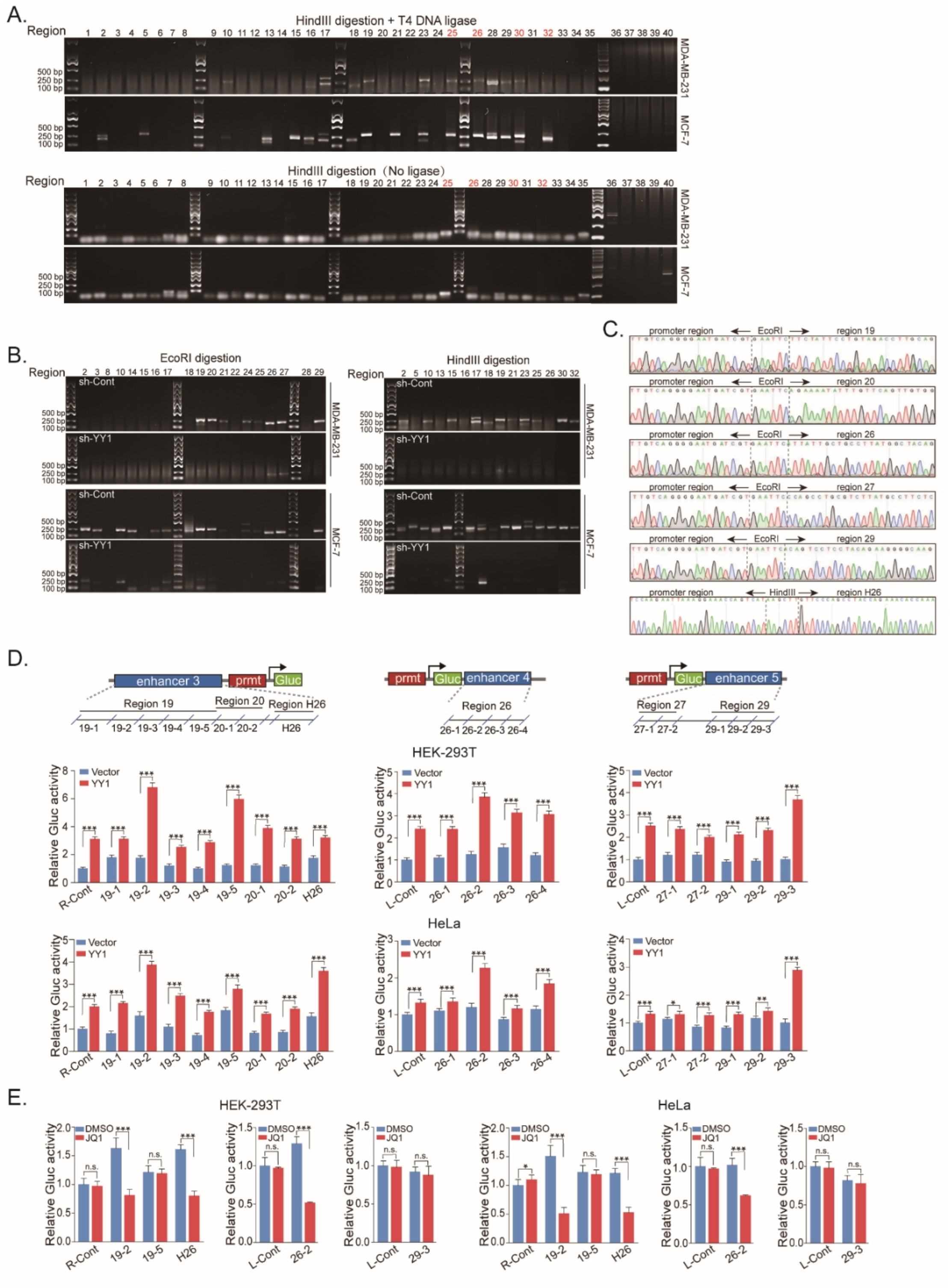

**Figure S6: YY1 recruits distal enhancers to activate FOXM1 gene expression.**

(**A**) Chromatin conformation capture (3C) assays to examine direct physical interactions between the FOXM1 promoter and its potential enhancer elements. The 3C assays followed the protocol described in Materials and Methods section, using HindIII to digest the genomic DNAs of MDA-MB-231 and MCF-7 cells. The ligation step was carried out in the presence or absence of T4 DNA ligase (top and bottom rows, respectively). The numbers on the top of the gels correspond to those in "EcoRI digestion site" in Figure 7A. PCR amplifications to examine the direct physical interactions between the FOXM1 promoter and each of the HindIII digested fragments used the primer sets shown in Table S4. The numbers of fragments overlapping with the predicted enhancer regions and showing interactions with the FOXM1 promoter are in red text. (**B**) The 3C assays carried out as described in A in MDA-MB-231 and MCF-7 cells infected by lentivirus expressing sh-Cont or sh-YY1. The genomic DNA fragments were generated by digestion of EcoRI (left panel) or HindIII (right panel). (**C**) DNA sequencing analyses to verify ligated bands in 3C assays. Ligations between the FOXM1 promoter and enhancer fragments digested by EcoRI or HindIII are indicated on the top. (**D**) Reporter assays to examine the effects of predicted enhancers on the FOXM1 promoter. Sectionalized enhancers 3, 4 or 5 were individually and ectopically placed adjacent to the FOXM1 promoter to generate different reporter constructs, as schematically illustrated in the top rows. Reporter constructs were transfected into HEK-293T and HeLa cells with or without ectopic YY1 expression to evaluate the effects of these sectionalized enhancer fragments and their response to YY1 expression (bottom rows). Data are shown as mean ± s.d. (n = 3). (**E**) Reporter assays to assess the effects of JQ1 on potential enhancer fragments of the FOXM1 promoter. The reporter constructs containing potentially active enhancer fragments adjacent to the FOXM1 promoter were transfected into HEK-293T and HeLa cells with or without JQ1 treatment, followed by examination of Gluc activity to verify the enhancer identity. Data are shown as mean ± s.d. (n=3).

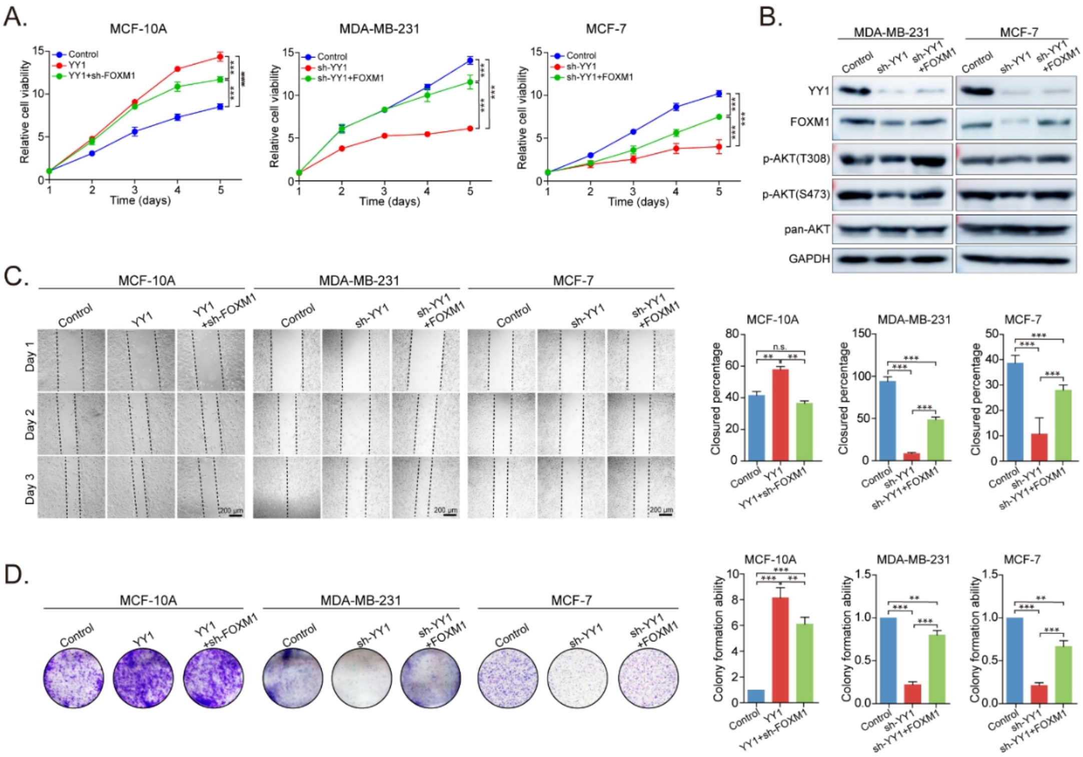


**Figure S7： YY1 promotes mammary cell proliferative transformation through FOXM1.** (**A**) Inverted FOXM1 expression attenuated the activity of YY1 in promoting mammary cell viability. MCF-10A cells expressing ectopic YY1 without or with FOXM1 knockdown by sh-FOXM1 were generated, with all shRNAs or genes delivered by lentiviruses. MDA-MB-231 cells carrying DOX-inducible sh-YY1 were individually infected by lentivirus expressing FOXM1 or an empty vector, and then cultured in medium containing 0.5 μg/ml of DOX to silence endogenous YY1. MCF-7 cells carrying sh-YY1 were individually infected by lentivirus expressing FOXM1 or an empty vector. Cell proliferation was determined by WST-1 assays, followed by calculation of cell viability. (**B**) Levels of AKT phosphorylation at S473 and T308, and total AKT in stably lentivirus infected MDA-MB-231 and MCF-7 cells with indicated genes or shRNAs. In Western blot analyses, the indicated antibodies were used to detect the levels of AKT phosphorylation and expression of different proteins. (**C**) Effects of manipulated YY1 and FOXM1 expression as indicated on the migration of mammary cells in wound healing assays. (**D**) Effects of manipulated YY1 and FOXM1 expression as indicated on the clonogenicity of mammary cells in colony formation assays.

**Supplementary Tables**

**Supplementary Table S1. Primers used in ChIP assays.**

| Name | Sequence (5' to 3') | Size (bps) |
|---|---|---|
| FOXM1-S1-For | ACGGTGTCTCCCTCTGTCG | 106 |
| FOXM1-S1-Rev | TCTGGAGGCTGAGGTGGAAGG | |
| FOXM1-S2-For | TGGGATTACAGGCGTGAGC | 109 |
| FOXM1-S2-Rev | GTTCTTTCCTTCCCCTGGGCT | |
| FOXM1-S3-For | GAACCTTGTCTGCCATTGTAT | 113 |
| FOXM1-S3-Rev | GGATATAATAGACCGTAGTGCT | |
| FOXM1-S4/5-For | TTCGGGAGGGGCAAAAGACAG | 103 |
| FOXM1-S4/5-Rev | CTTTGAGGGCTGCGTATTAT | |
| FOXM1-Neg-For | GGGAAGCGGGGGCGTGTCG | 116 |
| FOXM1-Neg-Rev | GGTGGGCGAGCCGAGGGAGAG | |

**Supplementary Table S2. Sequences of Cy5-labeled DNA probes, and wide type or mutant competitors.**

| Reported or predicted YY1 binding sites and their mutants | Names of probes or competitors | Sequences (5' to 3') |
|---|---|---|
| CDC6-S | Cy5-CDC6-For | CGCTCCCCGGCCATCTTGGCGGCTGGT |
| | Cy5-CDC6-Rev | ACCAGCCGCCAAGATGGCCGGGGAGCG |
| CDC6-SM | Cy5-CDC6M-For | CGCTCCGCGATTATCTTGGCGGCTGGT |
| | Cy5-CDC6M-Rev | ACCAGCCGCCAAGATAATCGCGGAGCG |
| S1 | S1-For | ATGATAAGCCATGATCACAC |
| | S1-Rev | GTGTGATCATGGCTTATCAT |
| S2 | S2-For | GGGCACAGACATTTTAATCA |
| | S2-Rev | TGATTAAAATGTCTGTGCCC |
| | Cy5-S2-For | GGGCACAGACATTTTAATCA |
| S3 | S3-For | CTTGTCTGCCATTGTATCTT |
| | S3-Rev | AAGATACAATGGCAGACAAG |
| S4 | S4-For | TGGTGCCGACATTTTTTTTC |
| | S4-Rev | GAAAAAAAATGTCGGCACCA |
| | Cy5-S4-For | TGGTGCCGACATTTTTTTTC |
| S5 | S5-For | GCTTTCTTCCATCTTGAAAA |
| | S5-Rev | TTTTCAAGATGGAAGAAAGC |
| | Cy5-S5-For | GCTTTCTTCCATCTTGAAAA |
| S2M | S2M-For | GGGCACAGAGGGTTTAATCA |
| | S2M-Rev | TGATTAAACCCTCTGTGCCC |
| | Cy5-S2M-For | GGGCACAGAGGGTTTAATCA |
| S4M | S4M-For | TGGTGCCGAGGGTTTTTTTC |
| | S4M-Rev | GAAAAAAACCCTCGGCACCA |
| | Cy5-S4M-For | TGGTGCCGAGGGTTTTTTTC |
| S5M | S5M-For | GCTTTCTTCGGGCTTGAAAA |
| | S5M-Rev | TTTTCAAGCCCGAAGAAAGC |
| | Cy5-S5M-For | GCTTTCTTCGGGCTTGAAAA |

**Supplementary Table S3. Primers used in the 3C experiments with EcoRI digestion.** The 23R primer is located in EcoRI digested fragments containing a FOXM1 promoter region, and all F primers are located in adjacent EcoRI digested fragments. PCR products were amplified using the primer pairs (23R primer and each of F primers).

| DNA Fragment | Type | Genomic sequence (5' to 3') | EcoRI position start | EcoRI position end | Size (bps) |
|---|---|---|---|---|---|
| FOXM1 locus (chr12; hg19) | | | | | |
| 1 | F | GGCAGCCCTGAGCACCACT | 3394853 | 3391090 | 246 |
| 2 | F | GATGCAGCCACACGGGTCTCC | 3391095 | 3380683 | 242 |
| 3 | F | TTAACTTCTCTGAGCCGCCCTATG | 3358351 | 3352504 | 215 |
| 4 | F | TTCTTTGTCTCTGTTCCACTGTCA | 3340476 | 3334132 | 272 |
| 5 | F | GGAGGAAGCAGCACTAAAAGCACT | 3326780 | 3320725 | 256 |
| 6 | F | GAGGGGAAGAAGAAAAGTGAATAA | 3308115 | 3304444 | 253 |
| 7 | F | GCAAGAACTAAGCACATCACAAAT | 3277634 | 3275328 | 256 |
| 8 | F | AATGAGGAATGAGCAGGACAGGAT | 3258273 | 3254754 | 244 |
| 9 | F | TGAATGAGCACGCCAAGGTT | 3236167 | 3229352 | 252 |
| 10 | F | GGCTGCTGACAAACAAAAGAAGGT | 3198457 | 3192759 | 231 |
| 11 | F | CTTCTCATCATTACTGTCATCAAA | 3191157 | 3185360 | 258 |
| 12 | F | CCATGATTATACCACTGCTCTCCA | 3185365 | 3183497 | 231 |
| 13 | F | GCATCCCCAGCAGACAGAACAGTG | 3168304 | 3165136 | 237 |
| 14 | F | GCTCACGCCTATAATCCTACCACT | 3125867 | 3120618 | 201 |
| 15 | F | GGAGGCCAGGATGTCAGGAG | 3099171 | 3092129 | 234 |
| 16 | F | GAGAAAACAAAAATCGCTAACCTT | 3084060 | 3066143 | 204 |
| 17 | F | TGCTTTCTCTTACTCAACAACTCT | 3045714 | 3044604 | 220 |
| 18 | F | GACCGGCGCAGTGGCTCAT | 3039507 | 3025388 | 214 |
| 19 | F | CTATCGTCACACCGCCACACTCC | 3018116 | 3009831 | 262 |
| 20 | F | AAGTCAATCCCGCATCCCGTTTTT | 3004605 | 3001517 | 273 |
| 21 | F | AATCAGTCAGACCATCCCGAATCA | 3001522 | 3000604 | 232 |
| 22 | F | GAAGTGGTGGTTGCAGTAAGGTGA | 3000609 | 2997678 | 242 |
| 23 | R | TGGATGGGAGGTAGAAGTGTCAAG | 2988094 | 2981211 | |
| 24 | F | CTAAGAGGAGGGCAGCAGTGACCA | 2952439 | 2936824 | 222 |
| 25 | F | ACATTTTATTATCCGGTGACTATT | 2926033 | 2922630 | 217 |
| 26 | F | TTGGCAAACAGCTGGGGAACT | 2922635 | 2915450 | 184 |
| 27 | F | GGGAAGCCAGGATGAGGAC | 2909230 | 2905289 | 199 |
| 28 | F | CAGAGGGTCAAATAGCCAAAGGTT | 2905294 | 2899184 | 293 |
| 29 | F | TTTCTAGCTGTGGACCCTTGAGTA | 2885237 | 2880430 | 209 |
| 30 | F | AGGCAGGGGAAGAGAAAGTGTAGC | 2862940 | 2861054 | 254 |
| 31 | F | TTCGCAATAGCCCAAAGGTAGAGA | 2846648 | 2844135 | 250 |
| 32 | F | CAGGGATATTGGACGGTAGTTTTC | 2832029 | 2825984 | 232 |
| 33 | F | CCCCATGGAAAAGCGCAGTATTAG | 2812215 | 2808883 | 225 |
| 34 | F | CCGTTGCAGGTGTGAGAGGTGTAT | 2801948 | 2798557 | 204 |

| | | | | | |
|---|---|---|---|---|---|
| 35 | F | ATTGTTAGGCTCATGTAGTTTTAT | 2780791 | 2767008 | 244 |

**Supplementary Table S4. Primers used in the 3C experiments with HindIII digestion.** The 27R primer is located in HindIII digested fragments containing a FOXM1 promoter region. All F primers are located in adjacent HindIII digested fragments. PCR products were amplified using the primer pairs (23R primer and each of F primers).

| DNA Fragment | Type | Genomic sequence (5' to 3') | HindIII position start | HindIII position end | Size (bps) |
|---|---|---|---|---|---|
| **FOXM1 locus (chr12; hg19)** | | | | | |
| 1 | F | CGTGCCCTGTGGAGTGCTGTTAC | 3401893 | 3398771 | 236 |
| 2 | F | AGCTGCCTGGGTCCTTCACACT | 3386844 | 3384831 | 263 |
| 3 | F | GGAGGGAAGAGGAGGGAAGGAAGA | 3352764 | 3347129 | 218 |
| 4 | F | GCAGGGCTAGAGGTGATGTGTTGT | 3347134 | 3342377 | 287 |
| 5 | F | CCCAGGCATACACAGAAACACAGG | 3329705 | 3320324 | 275 |
| 6 | F | CCTGAAATCCGCACCCTATGAAAT | 3316519 | 3300473 | 269 |
| 7 | F | CTCCTCTAGCTCCACCACCTGTCT | 3279040 | 3274238 | 193 |
| 8 | F | TTAACAAATGCTCCCAAAGTCAGG | 3265229 | 3259005 | 241 |
| 9 | F | CCCCCAAAATACCTCACGCATCT | 3254038 | 3242170 | 273 |
| 10 | F | GGCCCCCAAGATCCTGTTCG | 3240726 | 3229318 | 235 |
| 11 | F | GGCCTTTACCCCATCCCTTTCTC | 3229323 | 3222305 | 210 |
| 12 | F | GGCAATTCTGTGGCTTCCCTTCT | 3208889 | 3202263 | 203 |
| 13 | F | GGGCAGTTAACTTGGGGTGTGAGG | 3199606 | 3192112 | 253 |
| 14 | F | CTCTCCTCCCCTCCCCATTACTTA | 3189994 | 3187610 | 290 |
| 15 | F | GTAAACTAGGATCATGCCACTGTA | 3187420 | 3185175 | 253 |
| 16 | F | GGGGCATGCTCTTCTTCACTCCTT | 3166268 | 3162195 | 264 |
| 17 | F | CCAGAGACCCAGCTAGAAGGACAA | 3146994 | 3139731 | 226 |
| 18 | F | TGCCAAGTTCAAGAAGCCAGTCAC | 3139736 | 3137207 | 201 |
| 19 | F | GCTGGAGAAGGAGGCTATTTGTGG | 3118858 | 3111982 | 290 |
| 20 | F | TGACTGTTTTGACTTTTCCCTCTT | 3096908 | 3093566 | 241 |
| 21 | F | TCTGAAGAGTTAAAGGCAAGGTGA | 3086802 | 3065331 | 285 |
| 22 | F | GAAAAGGAAATAAGGGAAGAAATG | 3052084 | 3043817 | 248 |
| 23 | F | TACCCTATGACCACACGCCCTGAG | 3042694 | 3033278 | 296 |
| 24 | F | CACTGTGCCTGGGAAGATTTTTGT | 3009679 | 3006534 | 248 |
| 25 | F | GACCTCTGCCGCTTCCCATTCT | 3006539 | 2999491 | 270 |
| 26 | F | ATTCACTGGGGCACTCTCGGATTA | 2999475 | 2997106 | 277 |
| 27 | R | TTTGAGCTAATCGACCTGAATGGA | 2987712 | 2983950 | |
| 28 | F | GGAGTGAAGACATGGAGGCAAAAA | 2946162 | 2941455 | 242 |
| 29 | F | TGAGGCAGACAGGACAGTGAGGTT | 2941460 | 2935453 | 250 |
| 30 | F | TTCTGGTTGTGTACTGTGGGATGG | 2932589 | 2919509 | 286 |
| 31 | F | CCATGGGACCTTAAACTCTGACTT | 2907218 | 2905615 | 256 |
| 32 | F | GCCTCGGCCTCCCAAAGTTCTC | 2905620 | 2902729 | 275 |
| 33 | F | AGGGATTTTTAGTGAGGGTGGAGT | 2887822 | 2883039 | 262 |

| | | | | | |
|---|---|---|---|---|---|
| 34 | F | AATGATGCTCTGTAGAAAAATAAG | 2883044 | 2878623 | 247 |
| 35 | F | CCATCCCCCAACCTCCATTCTTTA | 2868810 | 2864040 | 221 |
| 36 | F | TGCACCTAGAGAAACAAGAACAAA | 2844721 | 2839951 | 249 |
| 37 | F | CCTCCACTCTTTTTACTCTCATTC | 2829863 | 2825779 | 237 |
| 38 | F | TGTTTGTGGAGAGCAGAATAATGA | 2815948 | 2808146 | 244 |
| 39 | F | TGCCTCAGCCTCCCAAAGTG | 2790984 | 2788375 | 203 |
| 40 | F | AGGAGTATGCAGAGGAGGCTTAGG | 2769937 | 2764996 | 205 |

**Supplementary Videos**

**Supplementary Video 1: Fusion process of EGFP-YY1 droplets.**

**Supplementary Video 2: Fusion process of EGFP-EP300-IDR3 droplets.**

**Supplementary Video 3: Fusion process of EGFP-EP300-IDR5 droplets.**
